# Pharmacokinetics and biodistribution of extracellular vesicles administered intravenously and intranasally to *Macaca nemestrina*

**DOI:** 10.1101/2021.07.28.454192

**Authors:** Tom Driedonks, Linglei Jiang, Bess Carlson, Zheng Han, Guanshu Liu, Suzanne E. Queen, Erin N. Shirk, Olesia Gololobova, Lyle H. Nyberg, Gabriela Lima, Liliia Paniushkina, Marta Garcia-Contreras, Kayla Schonvisky, Natalie Castell, Mitchel Stover, Selena Guerrero-Martin, Riley Richardson, Barbara Smith, Vasiliki Mahairaki, Charles P. Lai, Jessica M. Izzi, Eric K. Hutchinson, Kelly A.M. Pate, Kenneth W. Witwer

**Author notes:** these authors contributed equally (Tom Driedonks, Linglei Jiang).

## Abstract

Extracellular vesicles (EVs) have great potential as novel drug carriers for the treatment of various diseases. These lipid bilayer vesicles are naturally abundant in mammalian tissues and circulation, can be loaded with therapeutic small molecule drugs, (si)RNA, proteins and CRISPR/Cas9, and may be engineered for retention by specific tissues. However, many questions remain on the optimal dosing, administration route, and pharmacokinetics of EVs. Previous studies have addressed biodistribution and pharmacokinetics in rodents, but little evidence is available from larger animals. Here, we investigated the pharmacokinetics and biodistribution of Expi293F-derived EVs labelled with a highly sensitive nanoluciferase reporter (palmGRET) in a non-human primate model *(Macaca nemestrina),* comparing intravenous (IV) and intranasal (IN) administration over a 125-fold dose range. We report that EVs administered IV had markedly longer circulation times in plasma than previously reported in mice, and were detectable in cerebrospinal fluid (CSF) after 30-60 minutes. Already after one minute following IV administration, we observed EV uptake by PBMCs, most notably B-cells. EVs were detected in liver and spleen within one hour of IV administration. None of the IN doses resulted in readily detectable EV levels in plasma, CSF, or organs, suggesting that IN delivery of EVs in large animals including humans may require reconsideration or pretreatment approaches. Furthermore, EV circulation times strongly decreased after repeated IV administration, possibly due to immune responses and with clear implications for xenogeneic EV-based therapeutics. We hope that our findings from this baseline study in macaques will help to inform future research and therapeutic development of EVs.

---

Extracellular vesicles (EVs) are nano-sized vesicles produced by most or all cell types in multicellular organisms. EVs from specific cell types may also be harnessed as treatments for a wide range of human diseases and conditions, including cancer, inflammatory diseases, and tissue damage^1–4^. Furthermore, EVs may be loaded with therapeutic entities such as small molecule drugs^5^, proteins^6^, (si)RNA^7,8^, and CRISPR/Cas9^9,10^. The EV lipid bilayer protects its cargo from degradation and reduces off-target effects compared with nonencapsulated therapeutics^11^. Moreover, EVs may be engineered for retention by specific sites in the body, including brain, through the display of cell-specific surface motifs, usually proteins or peptides^12–14^. Since EVs occur naturally in the bloodstream and tissues, EV administration is thought to be safe and has reportedly elicited few toxic or inflammatory effects^15,16^. Although EVs are thus thought to be promising novel therapeutic tools, many questions remain about dosing, administration route, and pharmacokinetics.

To date, most pre-clinical studies have addressed the biodistribution and pharmacokinetics of EVs using mouse models^17–23^. Most studies have reported that EVs accumulate rapidly in the liver and spleen^18,19,21,23^, and sometimes lung^22^ and kidneys^17^. Additionally, EVs were found to have short circulation times in mice^17,19,21–23^. It is not well understood how EV administration route affects circulation time and biodistribution of EVs, since only a few studies have directly compared administration routes. In one study, EVs were administered to mice by intravenous (IV), subcutaneous, and intraperitoneal routes^18^. Compared with IV administration, subcutaneous and intraperitoneal administration resulted in lower EV uptake in liver and spleen, and higher uptake in the gastrointestinal tract and pancreas. Another study reported that intranasal (IN) administration of EVs resulted in improved brain targeting compared with IV administration^20^. While these mouse studies provide invaluable information on the biodistribution and therapeutic effects of EVs, results obtained in rodents may have limited translatability to human physiology^24^. Specific therapeutic effects of EV have been tested in sheep^25^, and pigs^26^, but pharmacokinetics studies on EVs in larger animals are scarce^3,27^.

Here, we investigated the pharmacokinetics of EVs in a non-human primate (NHP) model, the pig-tailed macaque *(Macaca nemestrina).* Large animal models allow repeated sampling from the same animal, in addition to sampling of multiple biofluids at the same time, and at volumes that cannot be obtained from small rodents. NHP are also physiologically similar to humans and are the best and in some cases only models of human disease. For example, NHP are exceptionally valuable to achieve better understanding of human immunodeficiency virus (HIV) disease progression and treatment, including assessment of HIV cure strategies and central nervous system disease^28^. Indeed, the study reported here is a prerequisite to trials of EV-associated transcriptional activators as latency-reversal agents for human immunodeficiency virus (HIV).

We used two relatively novel EV reporters: palmGRET, which is a palmitoylated EGFP-Nanoluciferase fusion protein^22^, and MemGlow 700, a near-infrared self-quenching lipid dye^29^. PalmGRET enables highly sensitive detection of EVs by emission of bioluminescence in the presence of a furimazine substrate^22^. MemGlow dye emits fluorescence in the nearinfrared range, where autofluorescence is generally reduced. Furthermore, this dye has been previously used to track tumor EVs in live zebrafish^30^.

For route of delivery, we compared IV administration, the most widely used route for systemic drug delivery^3^, with IN administration, which has been reported to achieve EV cargo delivery to the rodent brain^20^. It has been speculated that IN-administered small particles are transported by olfactory receptor neurons, which connect the nasal cavity and olfactory bulb to the brain^31^. For each administration route, we assessed how different EV doses affect the half-life of EVs in plasma and cerebrospinal fluid (CSF) of pigtailed macaques. Additionally, we measured the uptake of EVs by different subsets of peripheral blood mononuclear cells (PBMCs) shortly after administration. Furthermore, we compared the biodistribution of EVs in different organs of both macaques and mice. We found that the administration route strongly affected EV pharmacokinetics and tissue distribution. Repeated IV administration of EVs resulted in an accelerate blood clearance (ABC) phenomenon, potentially but not necessarily *via* immune-mediated effects. To our knowledge, this is the first reported study on the pharmacokinetics of EVs in macaques, which we trust will inform future studies on therapeutic applications of EVs.

## Results

### Production, separation, and general characterization of labelled EVs

To study the pharmacokinetics and biodistribution of EVs in larger animals such as macaques, highly sensitive reporters are required that can be detected with a high signal-to-background ratio. Therefore, we used two state-of-the art EV reporters: the near-infrared, self-quenching membrane dye MemGlow 700^29^, which was previously used to track tumor EV in live zebrafish^30^, and the dual reporter protein palmGRET (palmitoylated EGFP-Nanoluciferase), which was previously used to track tumor EVs in mice^22^. We transiently expressed palmGRET in Expi293F suspension cells (**Supplementary Figure 1A**) and harvested the conditioned culture medium three days later. Cells and debris were removed from culture medium by centrifugation and filtration. The EV-containing culture medium was concentrated tenfold by tangential flow filtration (TFF), a technique which is increasingly used for volume reduction during large-scale processing of EVs^32,33^. The EV concentrate was subsequently labelled with MemGlow 700, concentrated further by ultrafiltration, and subjected to size exclusion chromatography (SEC) to separate EVs from free dye and non-EV-associated proteins (**Supplementary Figure 1B**).

To satisfy the MISEV criteria^34^, we extensively characterized individual SEC fractions and pools by microBCA, SDS-PAGE and Western blot. A small protein peak was observed in SEC fractions 1–4 (**Supplementary Figure 1C and D**) which was positive for EV markers CD63, CD9, and TSG101 but devoid of ER marker calnexin (**Figure 1A**), indicating that EVs were isolated with minimal contamination by other cellular material. Transmission electron microscopy (TEM) and nanoparticle tracking analysis (NTA) of pooled fractions 1-4 confirmed the presence of EVs with expected morphology and an average diameter (by NTA) of 122.6 nm (SD +/− 9.9 nm) (**Figure 1B** and **C**). We performed further EV characterization using a bead-capture flow cytometry assay^35^, which enables profiling of 37 common EV surface markers (**Figure 1D**). Beside the major tetraspanins CD9, CD63, and CD81, the surface markers CD29 (integrin beta 1), CD146 (MCAM), and CD326 (EpCAM) were detected at considerable levels. Such integrins and adhesion molecules may steer the organotropism of EVs^36^. The surface protein CD47, which may prolong EV circulation times *in vivo* by reducing uptake by macrophages^37^, was additionally detected by Western blot, albeit at low levels (**Figure 1E**).

**Figure 1.**
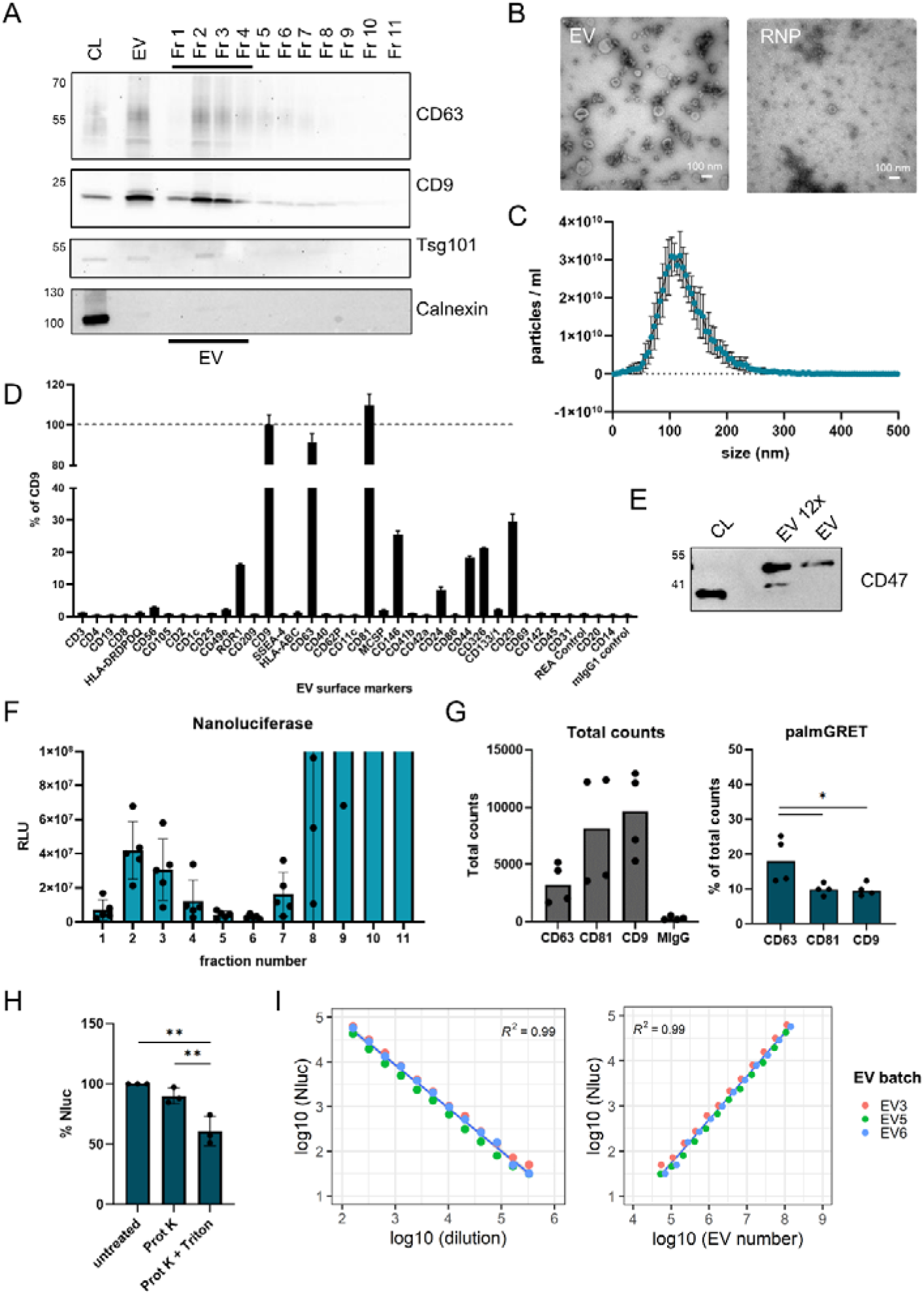
Characterization of palmGRET EVs. A) EVs produced by Expi293F cells were concentrated by TFF and ultrafiltration, followed by size-exclusion chromatography (SEC). Equal volumes of individual SEC fractions, pooled EV (fractions 1-4) and cell lysate were analyzed by SDS-PAGE and Western blot for EV markers CD63, CD9, TSG101, and ER marker *(i.e.* cytosol contaminant marker) Calnexin. Plot is representative of n=6 repeats. B) Pooled EV and protein fraction 10 (RNP) were imaged by negative stain TEM. Scale bars = 100 nm. Image is representative of n=6 batches. C) Particle size and concentration of pooled EV preparations were determined by NTA. Average of n=5 EV batches is shown. D) EV surface markers were profiled by MACSplex assay. Expression of surface markers is shown as percentage of CD9 expression. E) Western blot detection of CD47 in Expi293F cell lysates, TCA-precipitated EVs (12x), and EVs. F) SEC fractions 1–11 were diluted 20x, and the presence of the palmGRET reporter was validated by Nano-Glo assay. Measurement of n=5 EV batches is shown. G) SP-IRIS was used to determine the co-localization of palmGRET with CD63, CD81, and CD9. Total counts is the number of total fluorescent spots of n=4 EV batches that are captured by anti-tetraspanin and isotype antibodies. H) Pooled EVs were incubated with protease K and Triton X-100, protease K alone, or without additives. I) Pooled EVs were spiked into plasma of a healthy macaque, diluted in twofold serial dilutions from 160x — 327680x, and measured by Nano-Glo assay. Nano-Glo data were plotted against the dilution factor (left) and theoretical EV concentration from NTA (right).

### Characterization of label incorporation

We used a nanoluciferase assay to determine the presence of palmGRET in different SEC fractions (**Figure 1F**). Nanoluciferase was present in EVs (fractions 1-4) but was also highly abundant as free protein in the later protein fractions (7 – 11), highlighting the importance of size-based separation of EVs from non-EV proteins. An overview of the characteristics of different EV batches can be found in **Supplementary Table 1**. To confirm the incorporation of MemGlow 700, we used the Amnis Imagestream ISX imaging flow cytometer, which allows near-infrared detection and is suited to characterize small EVs^38,39^ (**Supplemental Figure 2**). To set gates for double-positive EVs, we used control EVs that contained only palmGRET or only MemGlow 700 (**Supplemental Figure 2A**). We performed serial dilutions to rule out coincidence events (**Supplemental Figure 2B**). Detergent treatment resulted in strongly reduced event counts, confirming that the measured events were indeed membrane particles (**Supplemental Figure 2C**) in accordance with the MIFlowCyt-EV recommendations^40^. On average, 30% of the EVs were double-positive for both reporters (**Supplemental Figure 2D**), while hardly any double-positive events were detected in unlabeled EVs, PBS, or free dye controls. Next, we used SP-IRIS to investigate the colocalization of palmGRET with the major EV tetraspanins CD9, CD63, and CD81 (**Figure 1E**). We observed the most palmGRET signal in EVs captured by anti-CD63 antibodies, and slightly less in EVs displaying CD81 and CD9, indicating that each of these tetraspanins was present in the palmGRET-labelled EV population. Next, we performed detergent/protease protection assays to confirm that palmGRET is enclosed within a lipid bilayer (**Figure 1F**). Protease K treatment of palmGRET EVs alone did not affect the nanoluciferase signal, whereas addition of detergent lead to a reduction in signal. This confirmed that palmGRET was enclosed within the lumen of EVs, as previously reported^22^.

**Figure 2.**
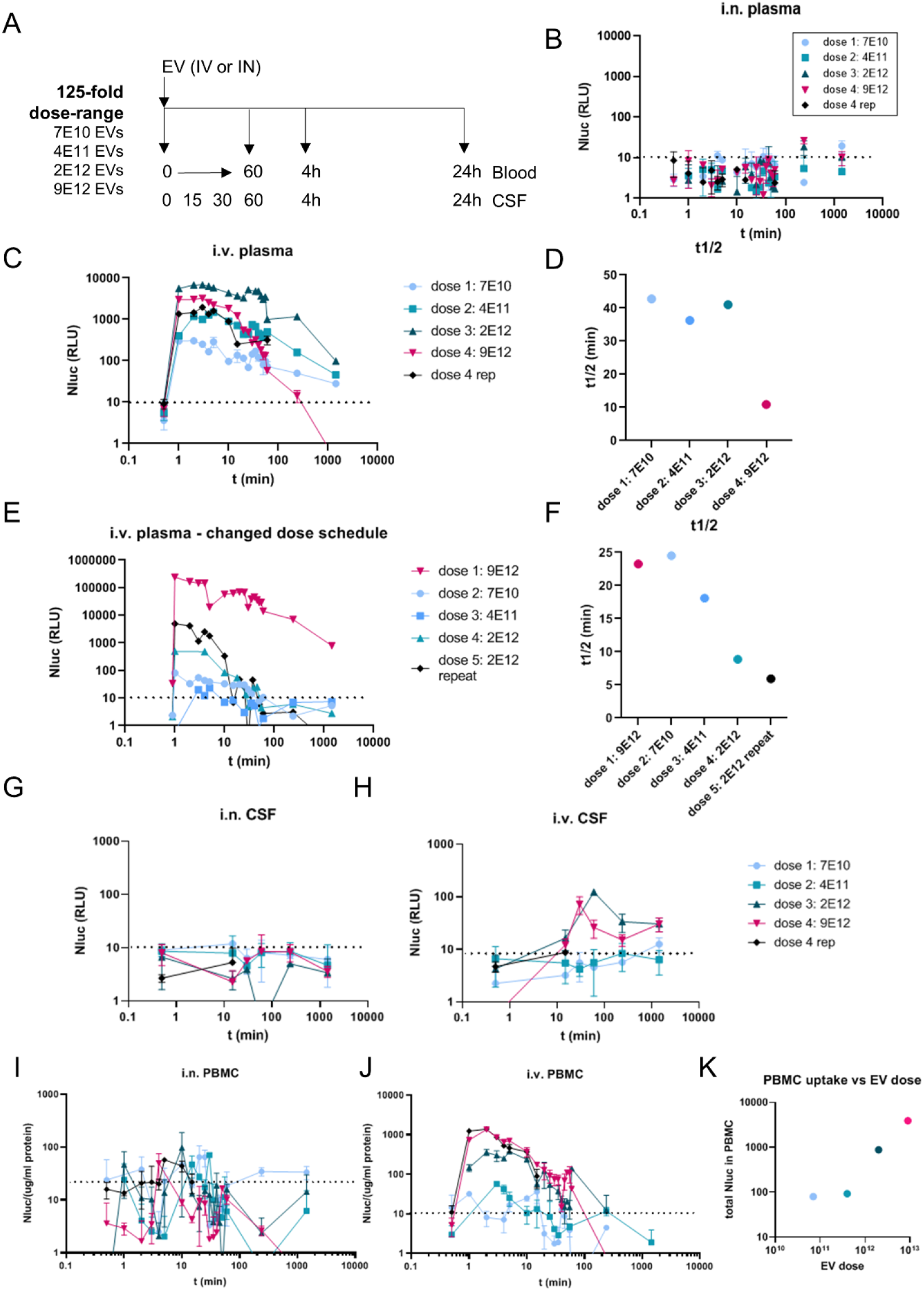
Pharmacokinetics of palmGRET EV administered intravenously and intranasally to macaques. A) Schematic of the study setup. EVs were administered intravenously (IV) or intranasally (IN) to macaques over a 125-fold dose range. Blood and CSF were sampled before administration, and at timepoints indicated in the chart. B) Detection of palmGRET EVs in plasma at different timepoints after intranasal administration to the first subject. C) Detection of palmGRET EVs in plasma after intravenous administration. D) EV half-life versus EV dose, from the data in Figure 2C. Nluc was plotted on a log-axis and t on a linear axis. The half-life was calculated from the slope between t=2 and t=60, using the formula t1/2 = log(2)/slope. E) Detection of palmGRET EVs in plasma after IV administration to a second subject with a different dosing schedule. F) EV half-life for each of the i.v. EV doses in Figure 2E, calculated as for Figure 2D. G) Detection of palmGRET EVs in CSF after intranasal administration to the second subject. H) Detection of palmGRET EVs in CSF after intravenous administration to the second subject. I, J) Detection of palmGRET EVs in PBMC lysates at different timepoints after IN (I) and IV (J) administration. K) Total nanoluciferase signal detected in PBMC lysates during the first 60 minutes after administration was calculated from Figure 2H, and plotted against the EV dose. palmGRET EVs were detected by Nano-Glo assay throughout. Error bars indicate standard error of the assay.

### Stability and detection in blood plasma

Prior to *in vivo* studies, we sought to determine detectability of labelled EVs in the biological matrix of blood plasma. This was done not only to assess assay sensitivity, but also because factors in blood might reduce stability of signal or contribute to background. palmGRET EVs were spiked into plasma, and serial dilutions were prepared over a 100,000-fold range (**Figure 1G**). Nluc signal was detected above background levels (macaque plasma without EVs) over the entire dilution range. We compared the Nluc signal with the number of particles per well (calculated from NTA particle counts and dilution factor), which suggested that the limit of detection was approximately around 200 EVs/μl (detection limit: 10,000 EVs in 50 μl = 200 EV/μl). We additionally tested the stability of different doses of palmGRET EVs in macaque serum for up to 24 hours (**Supplemental Figure 2E**). More than 60% of the initial dose was detectable in serum after 24 hours, with 80% for the two highest doses, indicating that palmGRET EVs are relatively stable in serum and not rapidly degraded by serum factors. These findings supported further use of the model to evaluate palmGRET EV pharmacokinetics and biodistribution.

### *Design and dosing:* in vivo *study*

We next compared intravenous and intranasal administration of different amounts of EVs, tracking the abundance of EVs in plasma and CSF over 24 hours after each administration (**Figure 2A**). The starting dose was based on a previous study in which EVs from 4E7 mesenchymal stem cells were administered into sheep fetuses^25^.^25^. We used a comparable number of cells to produce EVs for our starting dose. Initial measurements showed that Expi293F cells produced about 1.7E3 EV/seeded cell under the culture conditions we used. This set the starting dose to approximately 7E10 EVs (4E7 cells x 1.7E3 EV/cell ≈ 7E10 EVs). Three subsequent EV doses were administered at 5-fold greater concentration each time, with several weeks between doses. The fourth and highest dose was then administered a second time.

### Detection of EVs in blood plasma

Intranasal administration of EVs resulted in little if any nanoluciferase detection in plasma at any timepoint and after any dose (**Figure 2B**). This suggested that Expi293F EVs might have remained in the nasal cavity, mucosa, or lungs after administration, and did not enter the bloodstream. In contrast, intravenously administered EVs could be reliably detected in plasma at all doses (**Figure 2C**). For the three lowest doses (7E10, 4E11 and 2E12 EVs), nanoluciferase signal could be detected in plasma up to 24h (1440 minutes) after administration. Unexpectedly, the fourth and highest dose (9E12 EVs, magenta data points) was cleared more rapidly from plasma than dose 3, and was hardly detectable above background after 4 hours (360 minutes). We doublechecked the particle concentration and nanoluciferase signal in this particular EV dose, to rule out any issues with storage or handling, but we did not observe any abnormalities that could explain the observed lower signal and increased clearance. When we repeated administration of this highest dose (black data points), we observed a comparable clearance pattern.

### Half-life of EVs in blood plasma

Next, we used these data to calculate the half-life of EVs in plasma. The data followed a biphasic decay profile on a log-lin chart, with rapid decay shortly after administration followed by slower decay at a later timepoint, described by a two-compartment pharmacokinetic model^41^. We calculated the half-life from the data points during the first 60 minutes, which corresponds to the rapid decay phase in the model (**Figure 2D**). We observed an EV half-life between 36 and 42 minutes for the three lowest doses. In contrast, the half-life of signal after administration of the highest dose was approximately 11 minutes. Because we traced the clearance of the repeat of the highest dose for 15 minutes, the half-life of repeat dose 4 could not be reliably determined.

### Accelerated blood clearance: magnitude or number of doses?

The accelerated clearance observed at the highest dose could be due to the dose itself or to the repeated EV administration. We thus administered EVs to a naïve macaque in a different order, starting with the highest dose (9E12 EVs), then followed by the lowest dose (7E10 EVs) and increasingly higher doses again (**Figure 2E**). As before, EVs were detected in plasma after one minute and were cleared over time. Some signal remained detectable in plasma 24 hours following the highest dose, but not the other doses. However, the highest dose, now administered first, was cleared the slowest, while accelerated clearance was observed starting with the third EV administration (**Figure 2F**). Half-life decreased to ~9 minutes for dose 4 (2E12 EVs), and further to ~6 minutes for a repeat administration of the same dose. These results suggest that repeated administration, rather than the absolute EV dose, contributed predominantly to accelerated clearance.

### Accelerated clearance and inflammatory responses

We next asked whether the immune system and inflammatory responses might be involved in the accelerated blood clearance phenomenon. Inflammatory cytokines/chemokines were measured in plasma collected at different timepoints after administration of 9E12 EVs, for both the first and second subjects. (**Supplemental Figure 3A and B**). We did not observe induction of key inflammatory cytokines such as IL-6, TNFα, and IL-1β after EV administration, consistent with earlier reports^16^. IL-8 was detected at all timepoints (including baseline) after repeated dosing, and was induced after 4h in the naïve macaque, albeit with a high standard deviation. Additionally, MCP-1 was detected but did not show a trend consistent with induction. These results appear to be inconsistent with an inflammatory response to EV administration. In addition, total IgG levels in plasma, collected 24 hours after each dose, did not show a clear trend across the different doses (**Supplemental Figure 3C**). Thus, while EVs are cleared more rapidly from plasma after repeated administration, inflammatory responses and total IgG elevation do not appear to be involved.

### Detection of EVs in CSF

In addition to EV clearance from plasma, we tracked the uptake of EVs into the CSF after intranasal or intravenous administration. After intranasal administration (**Figure 2G**), EVs could not be reliably detected above background in CSF at any timepoint and after any dose. In contrast, signal was observed in CSF after intravenous administration at the higher doses (**Figure 2H**). EV signal in CSF peaked after 30 minutes for dose 4 and at 60 minutes for dose 3, and remained detectable above background for up to 24 hours. Intravenous doses 1 and 2 did not lead to detectable CSF at any of the timepoints. Intravenously administered EVs may thus migrate from plasma into CSF, at least at higher doses.

### Half-life of EVs in CSF and detection in plasma after intrathecal injection

We next questioned whether EV half-life in CSF is comparable to that in plasma. To achieve initial levels of EVs in CSF that were similar to those administered into blood, we introduced 3.2E10 EVs directly into the CSF of a previously untreated subject *via* intrathecal injection, followed by collection of CSF and plasma at regular intervals (**Supplementary Figure 4**). When injected into an estimated 15 ml of CSF, the EV concentration would be largely comparable to that of 2E12 EVs into 500 ml plasma (in the ~1E9 EV/ml range). Strong nanoluciferase signal in CSF remained detectable up to 6 hours. Based on data from the first hour post-treatment, EV half-life in CSF was approximately 12.5 minutes, considerably shorter than half-life in plasma. Meanwhile, nanoluciferase signal was detected above background in plasma only at later timepoints, suggesting that EVs might be able to diffuse from CSF into plasma, but at relatively low levels. Organs including brain were harvested 24 hours after administration, but nanoluciferase was not detectable at this late timepoint.

### Uptake of EVs by PBMCs

Since circulating white blood cells contribute to clearance of EVs from blood^4242^, we investigated uptake of EV-associated signal by peripheral blood mononuclear cells (PBMCs) at different timepoints following EV administration. Specifically, we isolated PBMCs from the blood samples collected in the 24h after administration, lysed them, and measured the amount of nanoluciferase taken up by these cells. After intranasal administration (**Figure 2I**), no EV uptake could be detected in the PBMCs at any of the doses, consistent with the absence of EV in blood plasma. In contrast, intravenous administration at all doses led to EV uptake as soon as one minute after injection (**Figure 2J**). At the highest doses, nanoluciferase was detectable in PBMCs up to one hour. There was also a linear relationship between total nanoluciferase signal (all time points combined) and dose for the highest three doses (**Figure 2K**). The repeat dose 4 was not included since blood samples could be collected over only 15 minutes.

### PBMC subtypes responsible for EV uptake

We next sought to determine the PBMC subtype(s) responsible for rapid uptake of EVs in blood. Flow cytometry was performed with PBMCs from whole blood samples collected during the first 10 minutes after administration of the highest EV dose (**Figure 3A**), using an antibody panel that identifies several PBMC subtypes: monocytes (CD159-CD3- CD20- and CD14+ or CD14-), T cells (CD3+ and CD4+ or CD8+), B cells (CD3-CD20+), and NK cells (CD159+). Granulocytes were gated based on their unique forward scatter/side scatter properties. EVs were detected based on the internal GFP label as well as by presence of the MemGlow self-quenching lipid dye. The gating strategy is depicted in **Supplementary Figure 5A**. We observed EV uptake by granulocytes, monocytes, CD3+ lymphocytes and CD20+ B cells already at one minute after administration (**Figure 3A**), consistent with nanoluciferase results from PBMC lysates (**Figure 2H**). Monocytes, CD3+ cells, and B cells differed in EV uptake efficiency: 80.8% of B cells became GFP+, while 14.1% of granulocytes, 13.8% of monocytes and 6.7% of CD3+ cells became GFP+. Of the CD3+ cells, both CD4+ and CD8+ T cells became GFP-positive to a similar extent (**Supplementary Figure 5B**). NK-cells did not efficiently take up EVs (**Supplementary Figure 5C**). We found similar percentages of EV uptake after repeating administration of the highest EV dose (**Figure 3B**). Gating of GFP+ or MemGlow+ cells showed largely similar uptake kinetics for both EV markers. However, B cells efficiently took up both GFP-containing and MemGlow-containing EVs, while CD3+ cells seemed to be less associated with the MemGlow signal. Granulocytes became GFP+ in the first 5 minutes, and MemGlow+ after 10 minutes. Since monocytes were less abundant than B cells and T cells, the percentage of MemGlow-positive monocytes could not be reliably determined. Taking into account both uptake efficiency and cell type contribution to the overall PBMC population, B cells were the largest positive population (~7% of all GFP+ mononuclear cells were B cells), followed by T cells (~4%), while monocytes were less than 1%. Overall, 11% of mononuclear cells took up GFP-containing EVs (**Figure 3C**).

**Figure 3.**
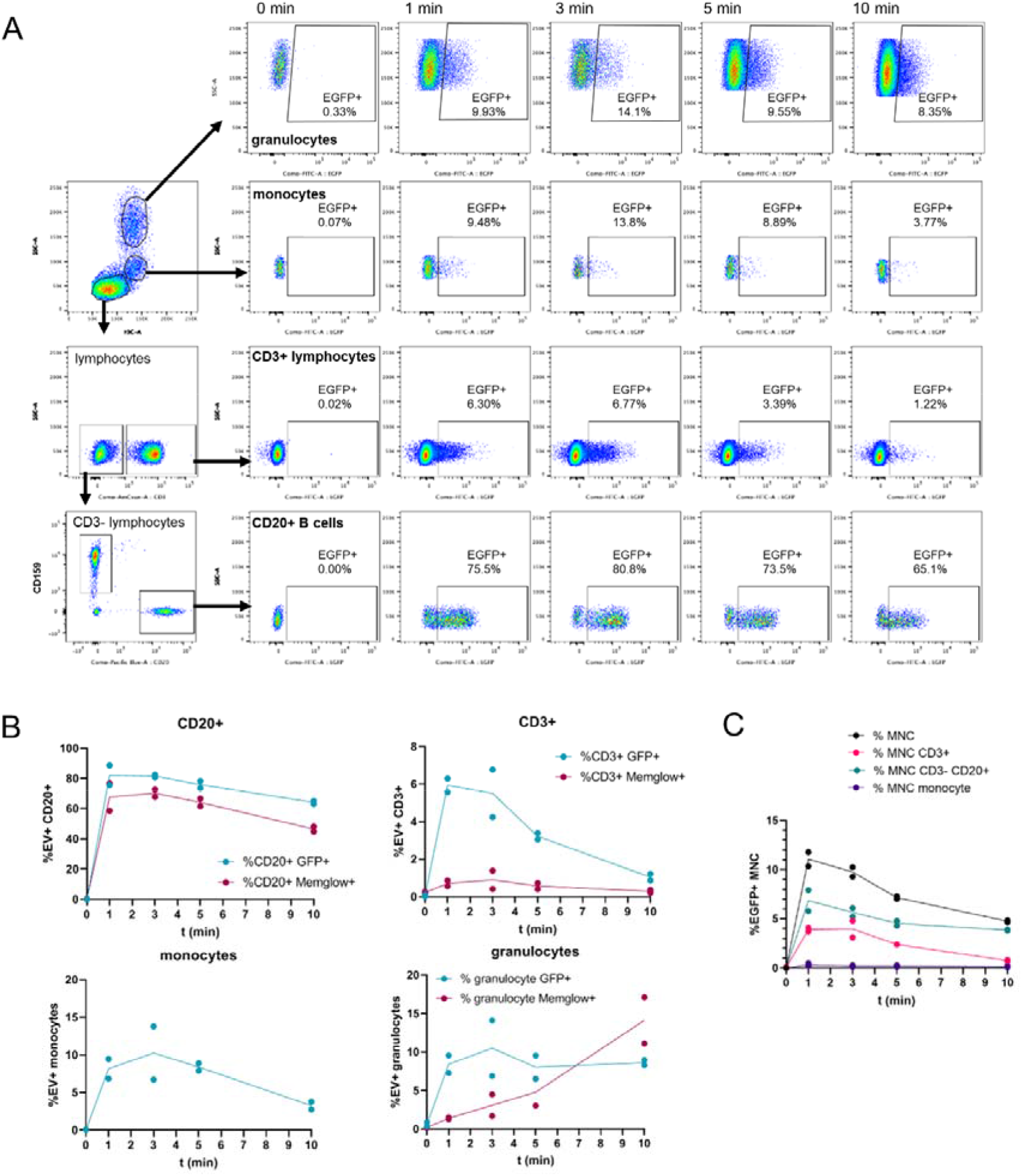
EV uptake by PBMC subsets quantified by flow cytometry. A) Whole blood collected after intravenous administration of the highest dose (dose 4) was immunolabeled, after which uptake of palmGRET (GFP)-containing EVs was measured by flow cytometry. The full gating strategy is found in Supplementary Figure 4A. EV uptake by granulocytes, monocytes, CD3+ lymphocytes and CD20+ B cells (bottom), presented as GFP+ cells as a percentage of each cell population. Plots are representative of n=2 EV administrations into the same animal, two weeks apart. B) Quantification of % GFP+ and % MemGlow+ PBMC subsets from Figure 3A, as percentage of each cell population. Data from n=2 EV administrations are shown. C) Quantification of GFP+ mononuclear cells (MNC), CD3+ lymphocytes, CD20+ B cells, and monocytes, relative to the total pool of MNC. Data from n=2 EV administrations are shown.

### Organ biodistribution in mouse

We also investigated how intranasal and intravenous administration affected distribution of our labelled EVs to different organs. For this part of the study, we performed initial experiments in mice, since their small size makes them suitable for *in vivo* and *ex vivo* imaging. For *in vivo* imaging, we used a modified nanoluciferase substrate that is more water-soluble than regular Nano-Glo, fluorofurimazine (FFz), allowing better distribution of the substrate throughout the whole animal^43^. After interperitoneal injection of FFz, we administered 1.4E11 EVs by intravenous or intranasal routes and measured bioluminescence (**Figure 4A**). After intranasal administration, we observed bright signal in the nasal cavity and in some cases also in the lungs. After intravenous administration, we observed most signal in the liver. After 40 minutes, we perfused the mice, harvested the organs and measured bioluminescence (**Figure 4B**) and near-infrared fluorescence *ex vivo* (**Figure 4C**). Most signal was observed in lungs (intranasal) or liver (intravenous), in line with our *in vivo* imaging observations. Intravenous administration additionally gave *ex vivo* bioluminescence in lung, spleen and kidney (**Figure 4B**), which was not observed in fluorescence mode (**Figure 4C**). Next, we prepared tissue homogenates from all harvested organs and measured EV uptake by nanoluciferase assay (**Figure 4D**). Intranasal administration did not result in strong nanoluciferase signal in most organs, although we observed high but variable nanoluciferase signal in lung, in line with our *in vivo* imaging results. Intravenous administration gave strong nanoluciferase signal in the liver and spleen, consistent with earlier reports^18,19,21–23^, and in kidney and lung, consistent with *ex vivo* bioluminescent imaging. Heart, colon and brain showed the lowest amount of nanoluciferase. EV uptake in brain was lower for intranasal administration than for intravenous administration. We also expressed the measured nanoluciferase signal as percentage of input dose per organ (**Supplemental Figure 6A**), and found that only a small percentage of the administered EVs was detected in the organs at the 40-minute timepoint.

**Figure 4.**
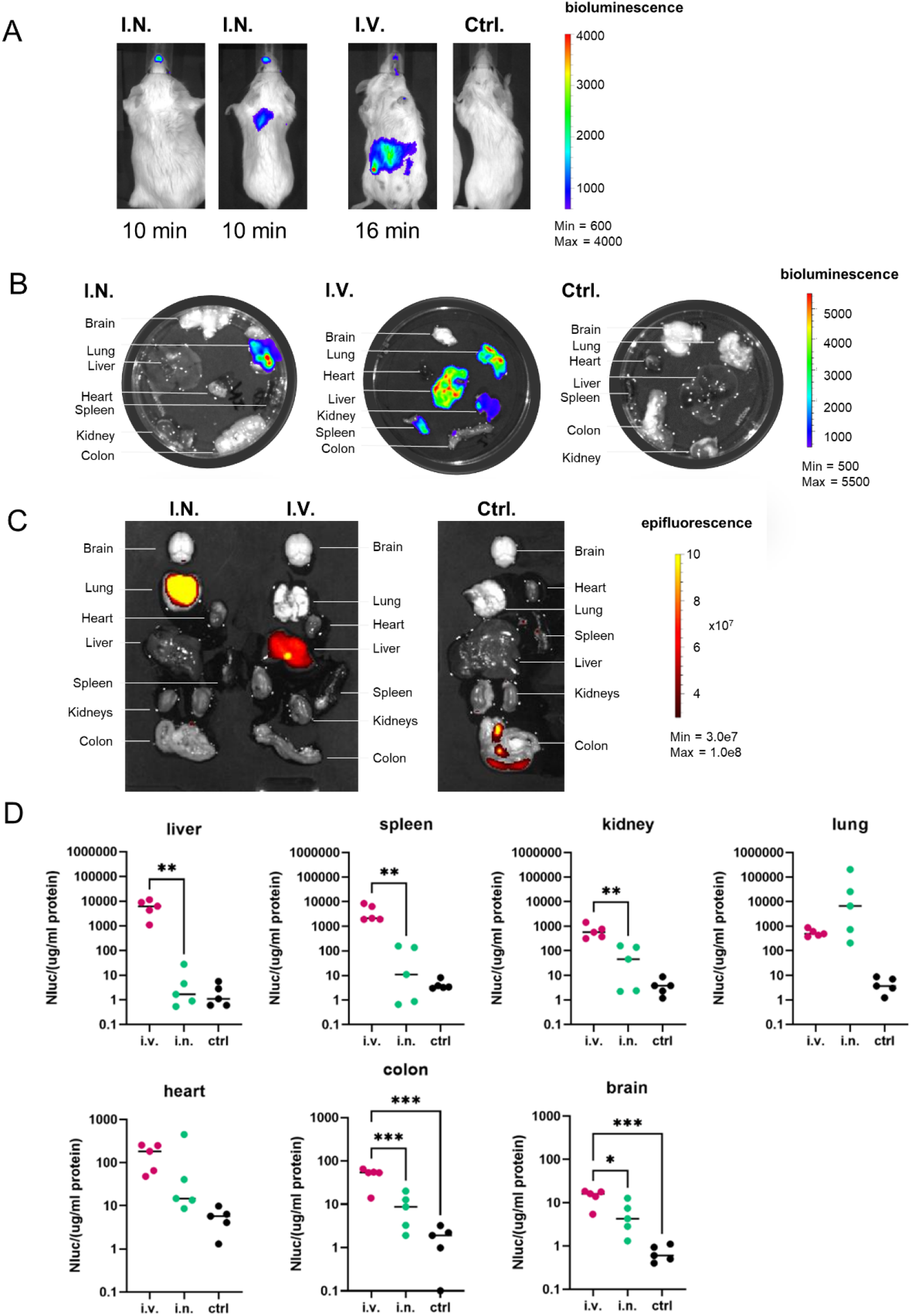
Biodistribution of palmGRET EVs administered intravenously or intranasally into mice. A) Female Balb/cJ mice were injected intraperitoneally with Nano-Glo substrate (fluorofurimazine, FFz), after which 1.4E11 EVs were administered intranasally (IN) or intravenously. *In vivo* biodistribution was monitored with a bioluminescence imager. Two intranasal administrations are shown (different outcomes), as well as one intravenous administration and one control, representative of n=2 experiments. B) *Ex vivo* imaging of mouse organs harvested 40 minutes after EV administration, in bioluminescence mode. C) *Ex vivo* imaging of mouse organs, in fluorescence mode (ex 698, em 713). Images are representative for n=2 experiments. D) EV uptake into mouse tissues was determined by Nano-Glo assay on tissue homogenates. Data from n=5 animals are shown. Statistical comparisons: one-way ANOVA with Tukey’s post-hoc test, * p < 0.05, ** p < 0.01, and *** p < 0.001.

### Organ biodistribution in macaque

EV biodistribution was also assessed by nanoluciferase assay of macaque tissues harvested 60 minutes after administration of the last, highest dose (9E12 EVs; **Figure 5A**) given to the first subject. After intravenous administration, we observed strong signal in liver and spleen, in line with the rodent results. Some uptake in lung was observed after intravenous, but not intranasal administration. EV uptake by kidney, heart, colon and brain was limited for both administration routes. We again calculated the nanoluciferase signal as percentage of the input dose per organ (**Supplemental Figure 6B**), and observed that, as in mice, only a small percentage of administered EVs could be detected in organs at the 60 minute timepoint. In addition to these peripheral organs, a faint nanoluciferase signal was observed in the medulla in brain after intravenous administration (**Figure 5A**, bottom right), suggesting this region might be the most accessible to EVs from the bloodstream. Nevertheless, the signal in medulla was low compared with signal in organs such as liver and spleen. Next, we used flow cytometry to identify which cell types in the spleen took up EVs (**Figure 5B**). After intravenous administration, we observed GFP+ monocytes and CD3+ lymphocytes, but the most efficient EV uptake was observed in B cells, consistent with the PBMC results reported above. Interestingly, after intranasal EV administration, we also observed a small percentage of GFP+ CD3+ lymphocytes in the spleen (**Figure 5C**).

**Figure 5.**
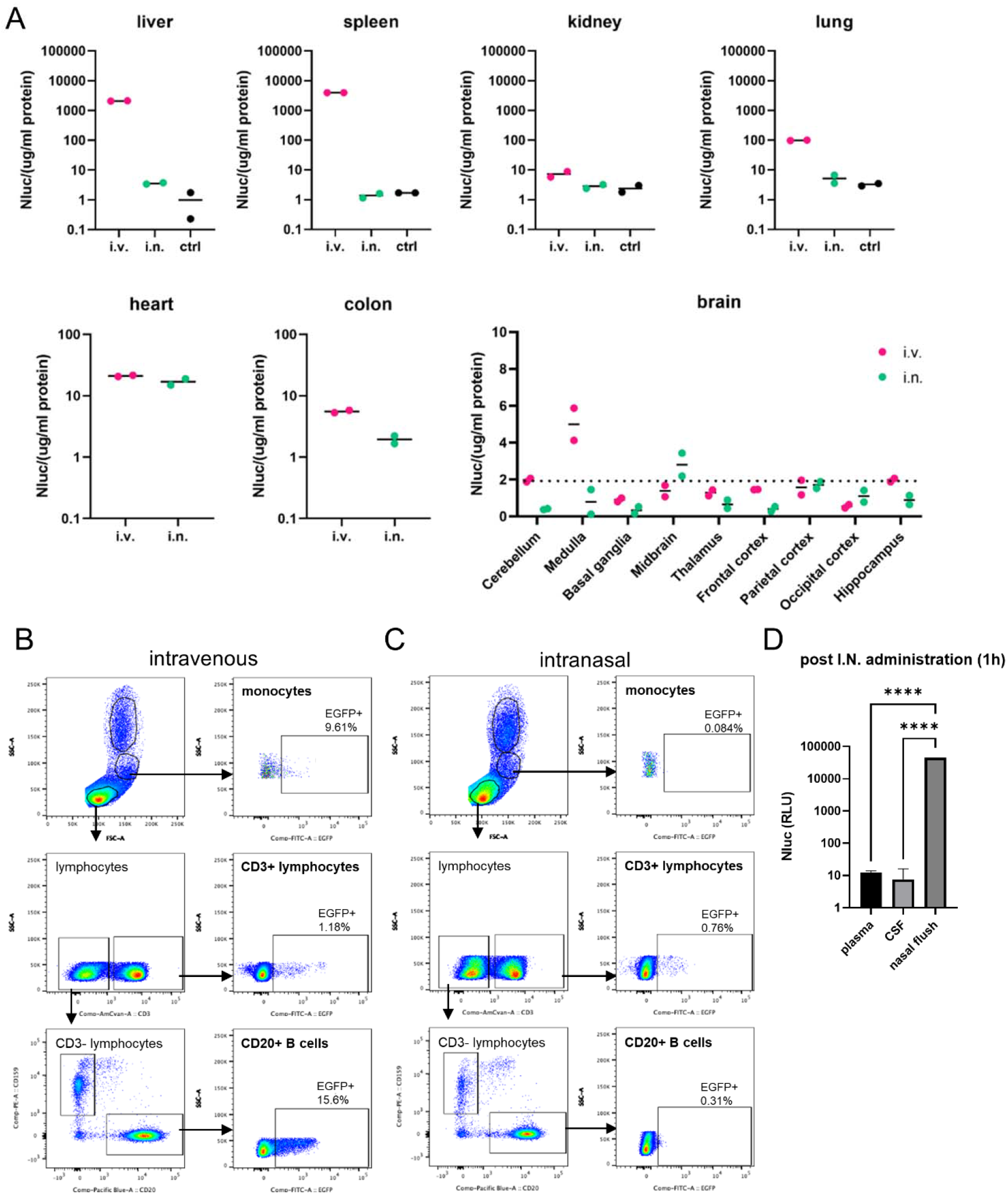
Biodistribution of palmGRET EVs administered intravenously or intranasally to macaques. A) Organs were collected 60 minutes after IV or IN administration of the highest dose of EVs (dose 4 repeat). EV uptake into macaque tissues was determined by Nano-Glo assay of tissue homogenates. Data from 1 animal per group is shown. Dots indicate replicate measurements of the same sample. B) Spleen was homogenized and immunostained directly for flow cytometric analysis of EV uptake by PBMCs in the spleen after intravenous administration. Gating strategy was the same as for whole blood PBMCs (see Supplementary Figure 4A). Dot plots show the % GFP+ cells as a percentage of the total B cells, CD3+ lymphocytes, or monocytes, respectively. C) Similar to B, EV uptake by PBMCs in the spleen after intranasal administration. Dot plots show the % GFP+ cells as percentage of the total B cells, CD3+ lymphocytes, or monocytes, respectively. D) The nasal cavity was flushed with PBS 1 hour after IN administration. Presence of EVs was measured by Nano-Glo assay, and compared with plasma and CSF of the same subject. **** p < 0.001 as determined by one-way ANOVA with Tukey’s post-hoc test.

### Barriers to intranasal uptake?

We next queried why intranasal administration did not result in systemic uptake of EVs. The *in vivo* bioluminescent images in mice showed a prominent signal in the nose of the animals, suggesting that EVs may be retained in the nasal cavity. To investigate this possibility in macaque, we administered 9E12 EVs intranasally into a naïve macaque. After 1 hour, the nasal cavity was lavaged with 10 ml PBS. We measured nanoluciferase activity in the nasal lavage fluid and observed a very strong signal compared with signal in plasma and CSF collected at the same time (**Figure 5D**), consistent with the nasal mucosa preventing EVs from being distributed to other locations.

## Discussion

### Half-life of EVs in plasma

To date, most preclinical studies on EVs have used mice, rats and zebrafish^3,27^ Remarkably, we measured the circulation time of EVs in NHP (t_1/2_ approx. 40 min) to be much longer than that reported in mice (t_1/2_ approx. 5 minutes)^17,19,21–23^ and zebrafish^42^. While most of these studies used different EV producer cells, which may have different clearance kinetics, one study reported that Expi293F EVs have a half-life of 10 minutes in mice^23^. In contrast, our EVs remained detectable in macaque plasma up to 24 hours after IV administration, suggesting that EV clearance may differ between different animal species. Comparing our results with the clearance of HIV-1 virions in macaques^44^, the half-life of our EVs was in the range of that of HIV-1 virions infused into naïve macaques, namely 13.0–19.3 minutes based on viral RNA, and 22–29 minutes based on pelletable Gag p24 in plasma^44^. Considering that HIV-1 actively fuses with and infects target cells, it is not surprising that EVs, which are thought to lack a consistent fusion mechanism like those of enveloped viruses, may have slightly longer circulation times. It has been suggested that species differences between the EV source (producer cell) and recipient animal model may affect circulation times. For example, human lentiviral vectors may be less stable in serum from evolutionary distant species^45^. Further study is required to determine if similar principles apply to the stability of EVs. However, we have shown *in vitro* that our EVs remained stable in macaque serum for at least 24 hours at 37°C.

### Accelerated clearance after repeated administration of EVs

An important consideration in potential EV therapeutics is whether repeated administration will provoke immune responses. If so, EVs might need to be prepared from autologous or allogeneic cells, as opposed to the much easier and cheaper alternatives of nonautologous or even xenogeneic materials^46,47^. In our study, EVs were cleared markedly more rapidly after the 4^th^ and 5^th^ administrations (repeated administration of the highest dose) than after administrations at lower doses and weeks earlier. By changing the order of the doses in follow-up experiments, we have shown that repeated dosing, and not the magnitude of the dose itself, led to accelerated clearance. We also did not observe a strong induction of inflammatory cytokines or total IgG levels, in line with previous findings in other models^12,15,16^. Accelerated clearance after repeat dosing is a known issue with PEGylated synthetic nanoparticles^19,48^, mediated by PEG-specific IgM^49^. It was recently shown that a similar principle applies to PEGylated EVs^50^. While EV-based vaccines can elicit strong immune responses^51–57^, these are designed to display or contain foreign antigens. Due to the species similarity between macaques and humans, we do not expect that EV-specific IgGs were elicited in our study. Furthermore, palmGRET EVs were relatively stable in macaque serum, arguing against a major contribution of the complement system. Various studies have shown that repeated administration of EVs gives strong therapeutic effects, such as a reduction in tumor burden^8,37,58^. The accelerated clearance that we observed is also not necessarily a roadblock to therapeutic applications of EVs, which, for many applications, must leave the circulation and enter tissues.

### Uptake by PBMC subtypes

Early after EV administration, we observed EV uptake into several PBMC subtypes. Our results differ somewhat from previous findings in that B cells appeared to take up EVs more efficiently than several other PBMC types. Once thought to be non-phagocytic, B cells are now known to have phagocytic capacity^59^. Indeed, a previous study also showed that human plasma EVs can be taken up by B cells^60^. However, in that study, the majority of uptake was observed in monocytes^60^. Previous *in vivo* work, *e.g.* with clodronate-depleted monocyte/macrophage populations^61^, and with blockade of scavenger receptor class A^62^, also implicated monocytes/macrophages in EV clearance. Possibly, the biological source of EVs, reporter system, species differences, or age of the subject contribute to these apparent differences. The rapid depletion of EV-associated signal from these cells is also worth noting, and could be due to degradation of EVs/label or to clearance of cells. Interestingly, within one hour of IV administration, we detected GFP+ B cells and monocytes in the spleen, suggesting rapid trafficking of blood B cells and monocytes to the spleen (and possible elsewhere) after EV uptake.

### Uptake into CSF, but not into brain

A striking result of our study was the low level of EV uptake into central nervous system compartments, regardless of administration route. First, intravenous administration: acceptance of the ability of EVs to cross the blood-brain barrier in both directions has become so widespread that statements to this effect are often not even referenced in the literature. Yet much of the evidence for EV transfer across the BBB is indirect. After intravenous administration, EVs might enter brain tissue directly across the BBB. They might also traverse the choroid plexus epithelium, a more permeable counterpart to the blood-brain barrier^63^.^63^. We detected low levels of EVs in brain tissue (especially medulla) and in CSF following IV administration, albeit at much lower levels than in peripheral compartments. In CSF, signal remained detectable at 24 hours. This result is similar to findings of a study of IV insulin in dogs, in which rapid clearance from plasma was followed by detection in CSF, peaking around 90 minutes^64^. However, we are not convinced that entry into CSF is an efficient precursor to brain entry. We did not observe any EV uptake into brain even after injecting EVs directly into the CSF (Driedonks and Witwer, data not shown). This suggests that reaching the CSF may not be enough to gain entry to the brain. Overall, the low levels of signal in both tissue and CSF suggest that the EV blood-brain route in our model is more of a precarious footpath than a superhighway.

### Negligible brain uptake after IN administration

Our results suggest that IN delivery of EVs to the brain in large animals should not be a foregone conclusion. Previously, studies of intranasally administered recombinant vesicular stomatitis viral vectors^65^ and a nanogel pneumococcal vaccine formulation^66^ in macaques found no brain uptake. Certainly, numerous studies report that IN EVs or their presumed cargo enter the brain parenchyma (see, for example^67–72^). As a result, intranasal delivery of EVs is thought to be a promising way to treat CNS disease (reviewed in^73^). However, the studies with positive results that we are aware of have all been performed in mouse or rat models, and predominantly with EVs sourced from MSCs or other stem cells, which seem to perform well in targeting the brain^73^. Physiology of the recipient species or characteristics of the source cells could explain disparate results. For example, many studies relied on various brain injury models (tumors, stroke, brain injury, morphine treatment) which may enhance the capacity to take up EVs compared with healthy animals^67,69,72^. Additionally, EV uptake and signal uptake may not overlap completely depending on EV separation technique. For example, one study finding efficient intranasal administration used MSC EVs that were incubated with gold nanoparticles and then ultracentrifuged for two hours at 100,000 g^20^. Although these procedures were meant to label EVs and remove free GNPs, it is unclear how efficient the labelling was and also doubtful that ultracentrifugation would separate EV-associated GNPs from free particles. Free GNPs might thus have contributed to or fully explained these results, with little or no EV uptake.

Intranasal delivery factors may influence outcome. Alternatives to instillation such as nebulization or aerosolization should be tested in large animals, in addition to instillation, and repeated short-term administrations might also be useful. Infusion of larger volumes leads to more entry into the lungs, instead of just the nasal cavity^74^, and could perhaps also influence brain uptake. Peptides or other adhesion molecules might be used on the EV surface to encourage uptake. One study used hyaluronidase treatment of the nasal cavity^70^, which may degrade extracellular matrix and enhance diffusion capacity^75^, remedying the nasal mucosa retention we observed. In any case, our findings suggest that different modes of intranasal delivery of different types of EVs should now be assessed in multiple models and perhaps with pre-treatments to determine if this route is a feasible EV delivery option for large animals including humans.

### EV uptake by peripheral organs

In both macaques and mice, IV-administered EVs were most efficiently taken up by the liver and spleen, followed by lung. This is in accordance with many other EV biodistribution studies in mice^18,19,21–23^. EVs were prominently detected in B cells in spleen after IV administration. It was recently reported that EVs may be taken up in the liver by Kupffer cells, hepatocytes and liver sinusoidal epithelial cells^76^, likely mediated by scavenger receptors^62^. However, we were unable to assess this in our macaque model. In mice, we observed low levels of uptake by kidney and, to a lesser extent, heart and colon. Thus, peripheral biodistribution of IV EVs was largely comparable between macaques and mice, with the exception of differences in kidney and lung. In mouse lungs, large EVs might be retained in the narrow microcapillaries (1-2 μm)^27^. Wider capillaries of macaques might explain the relatively reduced retention of EVs. It is important to note that EV biodistribution studies based on reporter proteins cannot show how many EVs cumulatively reached an organ, since protein-based reporters may be degraded over time^21^. Instead, these data can be used to reliably compare the relative uptake of EVs by different organs at a certain timepoint. Furthermore, we cannot account for EVs that have been taken up by endothelial cells, for example those of the vascular system.

### Factors beyond administration route

that affect EV biodistribution may include differences in tetraspanins or other surface proteins, EV labeling strategy^21,23^, and the type of EV donor cells^18,77^. Illustrating this, endogenous CD63-Nluc EVs from cardiomyocytes were shown to target different organs than CD63-Nluc EVs from Expi293F cells^23,77^. We have investigated the biodistribution of EVs from only one cell source, which may not be representative of EVs from, *e.g.,* MSCs or red blood cells. Additionally, the disease status of the recipient may affect tissue accumulation. Uptake of MSC-EVs into kidneys was increased in mice with acute kidney injury compared to healthy controls^78,79^. Future research will likely highlight other factors that control EV tissue retention (or even true “homing” or “targeting,” if this is possible for EVs^13^), and will give clues on how to better control the fate of EVs *in vivo.*

Taken together, nanoluciferase-based reporters allowed sensitive tracking of EVs in larger animals for pharmacokinetic measurements. We show that EVs from human cells had a longer circulation time in macaques compared with mice, but that CNS penetration was low for both IV and IN administration. Repeated administration led to more rapid clearance, which may have implications for EV-based therapies against cancer and immune diseases. We hope that our findings from this baseline study in macaques will help to inform future research and therapeutic development of EVs.

## Materials & Methods

### Cells and plasmids

Expi293F cells (Thermo Fisher) were maintained in Expi293 medium (Gibco, Waltham, MA) in vented shaker flasks on a shaker platform maintained at 125 rpm in a humidified 37°C incubator with 8% CO_2_. The pLenti-palmGRET reporter^22^ was provided by C.P. Lai (Addgene 158221), and endotoxin-free plasmid DNA megapreps were prepared by Genewiz (Genewiz, South Plainfield, NJ). For each EV production batch, 3 x 1L shaker flasks were seeded with a total of 750 ml of cell suspension at 3E6 cells/ml. Cells were transfected using Expifectamine (Thermo Fisher) according to the manufacturer’s instructions, with 150 μg pDNA (0.6 μg pDNA per ml of culture) and 480 μl of Expifectamine per flask. One day after transfection, 1.2 ml Enhancer 1 and 12 ml Enhancer 2 were added to each flask. Cultures were harvested three days after transfection. Transfection efficiency was checked on a Nikon Eclipse TE200 fluorescent microscope, cells were counted on a hemacytometer and tested for viability by Trypan blue exclusion (Thermo Fisher). Cell densities and viability of different batches at harvest are found in **Supplementary Table 1.**

### EV separation and fluorescent labelling

Cells were removed from conditioned medium by centrifugation at 1000 g for 20 min at 4C. Supernatant was centrifuged again at 2000 g for 20 min and filtered through a 0.22 μm bottle-top filter (Corning, NY). EVs were concentrated to 75 ml by tangential flow filtration (TFF) using two 100 kD Vivaflow 50R cassettes (Sartorius, Goettingen, Germany) run in parallel on a Cole-Parmer Masterflex L/S peristaltic pump operated at 100 rpm. The concentrated EVs were fluorescently labelled by adding 200 nM MemGlow 700nm dye^29^ (Cytoskeleton Inc., Denver, CO) and incubating at RT for 30 min. EVs were concentrated further on Amicon 15 Ultra RC 100kD filters (Millipore Sigma, Darmstadt, Germany), spun for 20 – 30 min at 4000 g. The concentrate was loaded onto a qEV10 70nm SEC column (Izon, Medford, MA) run with DPBS (Gibco, Waltham, MA), after discarding the void volume, 5-ml fractions were collected. EV-enriched fractions 1-4 were pooled together and were concentrated again on Amicon 15 Ultra RC 100kD filters, spun for 20 – 30 min at 4000 g. EVs were aliquoted and stored in LoBind tubes (Eppendorf, Bochum, Germany) at −80°C.

### Trichloroacetic acid (TCA) precipitation

For CD47 immunoblotting, EVs were precipitated using TCA. 230 μl Expi293F EVs were thawed from storage at −80°C, after which 2 μl of 2% sodium deoxycholate (Sigma Aldrich) was added. Tubes were vortexed and incubated at room temperature for 15 min. 23 μl of 100% TCA (Sigma Aldrich) was then added, samples were centrifuged at 14,000 rpm for 15 min at 4°C, and the supernatant was aspirated. Precipitated pellets were washed once with 500 μl ice-cold acetone and centrifuged again at 14,000 rpm for 15 min at 4°C. After removal of supernatant, samples were air-dried for 10 min and pellets dissolved in sample buffer for immunoblotting.

### Immunoblotting

Transfected Expi293F cell pellets were lysed in PBS + 1% Triton-X100 and Complete protease inhibitor tablets (Roche, Mannheim, Germany) for 15 minutes on ice. Nuclei were spun down for 15 minutes at 14,000 rpm in a tabletop centrifuge at 4°C. Cell lysate, final EV isolate and individual SEC fractions were mixed with 4x TGX sample buffer (Bio-Rad, Hercules, CA) under non-reducing conditions (except for CD47 blotting, under reducing conditions), boiled for 5 minutes at 100°C, and subjected to PAGE gel electrophoresis on a 4%-15% Criterion TGX Stain-Free Precast gel (Bio-Rad). Proteins were transferred to a PVDF membrane using the iBlot2 system (Thermo Fisher) run on program p0. After 1h of blocking in 5% Blotting-Grade Blocker (Bio-Rad) in PBS + 0.05% Tween-20 (PBS-T), blots were incubated overnight at 4°C with the following primary antibodies in blocking buffer: Rabbit-anti-Calnexin (1:1000, ab22595, Abcam), mouse-anti-CD63 (1:3000, #556016, BD Biosciences), mouse-anti-CD9 (1:3000, #312102, BioLegend), rabbit-anti-Tsg101 (1:2000, ab125011, Abcam), or rabbit-anti-human CD47 (1:500, Cusabio Technology, CSB-PA005993). Blots were washed 3x with PBS-T and incubated for 1h at room temperature with secondary antibodies mouse-IgGk-BP-HRP (sc-516102, SantaCruz) or mouse-anti-rabbit-IgG-HRP (sc-2357, SantaCruz) diluted 1:10,000 in blocking buffer. After washing 3x with PBS-T and 2x with PBS, SuperSignal West Pico PLUS Chemiluminescent Substrate (Pierce, Rockford, IL) (or, for CD47, SuperSignal West Femto Maximum Sensitivity) was used for detection on an iBright FL1000 (Thermo Fisher) in chemiluminescence mode.

### Nanoparticle tracking analysis

ZetaView QUATT-NTA Nanoparticle Tracking Video Microscope PMX-420 and BASIC NTA-Nanoparticle Tracking Video Microscope PMX-120 (ParticleMetrix, Inning am Ammersee, Germany) were used for particle quantification in scatter mode. The system was calibrated with 100 nm PS beads, diluted 1:250,000 before each run. Capture settings were sensitivity 75, shutter 100, minimum trace length 15, cell temperature was maintained at 25°C for all measurements. Samples were diluted in 0.22 μm filtered PBS to a final volume of 1 ml. Samples were measured by scanning 11 positions twice, recording at 30 frames per second. Between samples, the system was washed with PBS until no particles remained. ZetaView Software 8.5.10 was used to analyze the recorded videos with the following settings: minimum brightness 20, maximum brightness 255, minimum area 5, and maximum area 1000.

### Transmission electron microscopy

10 μL sample was adsorbed to glow-discharged carbon-coated 400 mesh copper grids by flotation for 2 minutes. Grids were quickly blotted and rinsed by flotation on 3 drops (1 minute each) of Tris-buffered saline. Grids were negatively stained in 2 consecutive drops of 1% uranyl acetate (UAT) with tylose (1% UAT in deionized water (diH2O), double filtered through a 0.22 μm filter), blotted, then quickly aspirated to cover the sample with a thin layer of stain. Grids were imaged on a Hitachi 7600 TEM operating at 80 kV with an AMT XR80 CCD (8 megapixel).

### Single Particle Interferometric Reflectance Imaging Sensing (SP-IRIS)

EVs diluted 1:100 in DPBS were diluted 1:1 in incubation buffer (IB) and incubated at room temperature on ExoView R100 (NanoView Biosciences, Brighton, MA) chips printed with anti-human CD81 (JS-81), anti-human CD63 (H5C6), anti-human CD9 (HI9a) and anti-mouse-IgG1 (MOPC-21). After incubation for 16 hours, chips were washed with IB 4 times for 3 minutes each under gentle horizontal agitation at 500 rpm. Chips were then incubated for 1h at RT with fluorescent antibodies anti-human CD81 (JS-81, CF555) and anti-human CD63 (H5C6, CF647) diluted 1:200 in a 1:1 mixture of IB and blocking buffer. Anti-human CD9 was not used, since the third wavelength was needed for the palmGRET reporter protein. The chips were subsequently washed once with IB, three times with wash buffer, and once with rinse buffer (all washes 3 minutes at 500 rpm agitation). Chips were immersed twice in rinse buffer for 5 seconds and removed at a 45° angle to remove the liquid from the chip. All reagents and antibodies were obtained from NanoView Biosciences (#EV-TETRA-C). All chips were imaged in the ExoView scanner (NanoView Biosciences) by interferometric reflectance imaging and fluorescent detection. Data were analyzed using ExoView Analyzer 3.0 software. Fluorescent cutoffs were as follows: CF555 channel 300 a.u., CF488 channel 410 a.u., CF647 channel 300 a.u., allowing <1% of particles above background in the isotype control. Fluorescent counts from multiple measurements were normalized against the total fluorescent particle count.

### Imaging flow cytometry

EVs diluted 1:10 in DPBS were quantified by imaging flow cytometry on an Amnis ImagestreamX MkII instrument (Amnis Corp, Seattle, WA) on low flow speed, using a 60x objective and extended depth of field (EDF) option enabled. EGFP signal was collected in channel 2 (480-560 nm filter), MemGlow 700nm signal was collected in channel 5 (642-745 nm filter), and sideward scatter (SSC) was collected in channel 6 (745-800 nm filter). Negative controls recommended by the MiFlowCyt-EV consortium^40^ were included in all measurements: buffer only control, free dye control (200 nM MemGlow700 in PBS), single stained EVs (palmGRET only, and MemGlow700 labelled EV from untransfected Expi293F cells). Serial dilutions were included to ensure measurement in the linear range of the instrument, and to rule out swarm effects. Data were analyzed using Amnis IDEAS software v6.2.

### MACSPlex surface marker characterization

EV surface markers were characterized *via* MACSPlex (Miltenyi Biotec, #130-122-209), following manufacturer’s instructions. In brief, 120 μl EV sample was mixed with 15 μl MACSPlex capture beads and incubated in a shaker at room temperature overnight. Next, 500 μl MACSPlex buffer was added, centrifuged at 3000 x g for five minutes, and 500 μl supernatant was removed. 15 μl MACSPlex detection reagent cocktail was added (anti-CD9, -CD63 and -CD81, as included in kit), and samples were incubated for one hour. After two washes in 500 μl assay buffer, data were collected on a BD LSR Fortessa flow cytometer. For flow cytometric data analysis, median signal intensity was used for relative quantification of surface markers. PBS (negative control) signal was subtracted from the median signal intensity of each sample. The median signal intensity of CD9 beads was used to normalize the abundance of other surface markers.

### *In vivo* administration

#### Mice

Balb/cJ mice (Jackson Labratories, 8-12 weeks, female) were injected intraperitoneally with 100 μl fluorofurimazine^43^ under sedation with isoflurane. Subsequently 1.5E11 EVs were administered by tail-vein injection or intranasal instillation. Bioluminescent imaging was performed on a Caliper IVIS SpectrumCT *In Vivo* Imaging System (Caliper Life Sciences, Hopkinton, MA, USA). Images were taken every 30 seconds (exposure time = 30 s) in luminescence mode. Mice were perfused with PBS *via* cardiac puncture 40 minutes after EV administration, organs were harvested and imaged *ex vivo* in fluorescence mode (ex 689, em 713), and in bioluminescence mode while immersed in NanoGlo substrate (diluted 1:50, Promega #N1110). Tissues were homogenized in 1 ml N-PER (brain) or T-PER (other organs) + Complete Mini protease inhibitor cocktail tablet (Roche, Mannheim, Germany) in FastPrep Lysing Matrix D tubes on a FastPrep homogenizer. Homogenates were centrifuged for 5 min at 10,000 g at 4°C, supernatant was taken off and used in NanoGlo and BCA assays (Pierce, Rockford, IL). Mice experiments were performed under approval of the Johns Hopkins University Animal Care and Use Committee (ACUC), study number M018M145.

#### Macaques

Juvenile pigtailed macaques (*Macaca nemestrina,* male, 3-4 years old) were obtained from the JHU pigtail colony. EVs (7E10 EVs, with 5-fold increments for all subsequent doses) were administered by intravenous injection into the small saphenous vein or by intranasal instillation through a catheter under ketamine sedation (10 mg/kg body weight). Macaques remained sedated during blood/CSF collection for the first hour by administering ketamine in 10-20mg increments, and were sedated again with 10 mg/kg body weight at the 4h and 24h timepoints. After each EV dose and biofluid collection, macaques were given two weeks to recover until the next EV administration, for a total of 5 doses. To assess the turnover rate of EVs in CSF, we administered 3E10 EVs *via* intrathecal injection into the subarachnoid space, and collected blood and CSF after 0, 0.5, 1, 3, 6, and 24 hours. To assess retention of EVs in the nasal cavity, nasal lavage was performed 1 hour after intranasal EV administration using 10 ml PBS, spun down at 2000 g for 10 min. Macaque experiments were performed under approval of the Johns Hopkins University Animal Care and Use Committee (ACUC).

At each timepoint, 500 μl blood was collected by venipuncture into tubes containing 100 μl ACD. Blood was processed within 1.5h after collection by centrifugation for 5 minutes at 800 g, plasma was taken off and stored directly at −80°C. To collect PBMC from the same sample, the blood cell pellet was reconstituted to 1 ml with DPBS, carefully layered onto a 1 ml Histopaque-1077 cushion (Sigma, St. Louis, MO), and centrifuged for 30 min at 400 g without brake. After discarding the supernatant layer, the PBMC-containing interphase was transferred to a new tube. PBMCs were washed twice by adding 1 ml PBS, centrifuging for 10 min at 250 g, and discarding the supernatant. The final PBMC pellet was taken up in 200 μl lysis buffer (PBS + 1% Triton-X100 + Complete Mini protease inhibitor cocktail tablet (Roche, Mannheim, Germany)), lysed on ice for 15 minutes. Nuclei were removed by centrifuging 15 minutes at 16,000 g at 4°C. Protein concentration was determined by BCA assay (Pierce, Rockford, IL). Per timepoint, 500 μl CSF was collected which was centrifuged for 10 minutes at 2000 g to remove cells. The supernatant was taken off and stored directly at −80°C. After the final dose, macaques were euthanized using Nembutal (20-30 mg/kg) and perfused with PBS. Organs were excised and snap-frozen at −80°C. Parts of the spleen and bronchial lymph nodes were processed directly for flow cytometry (see below for details).

### Flow cytometry

PBMCs were immunolabeled directly in whole blood with fluorescent antibodies. 100 μl whole blood was added to antibody cocktails (mo-anti-CD159a-PE, Beckman Coulter, cat# IM3291U, dil 1:30; mo-anti-CD4-PerCP/Cy5.5 BD Biosciences 552838 dil 1:7.5; mo-anti-CD20-e450 Thermo Fisher, cat# 48-0209-42 dil 1:60; mo-anti-CD3-V500 BD, cat# 560770, dil 1:30; mo-anti-CD8-BV570, BioLegend, cat# 301038, dil 1:60; mo-anti-CD14-BV650, BioLegend, 563419, dil 1:30), briefly vortexed and incubated at room temperature for 20 minutes. Next, red blood cells were lysed by addition of 2 ml RBC lysis buffer (ACK lysing buffer 0.83% NH_4_Cl, 0.1% KHCO_3_, 0.03% EDTA) and incubation at room temperature for 10 minutes. Tubes were centrifuged at 400 g for 5 minutes and supernatant was discarded. Next, 2 ml PBS was added and tubes were centrifuged again at 400 g for 5 minutes. Supernatant was discarded, labelled PBMCs were carefully resuspended in 500 μl PBS and measured directly on a BD LSR Fortessa flow cytometer. As negative controls, we included whole blood collected before the injection of EVs, and fluorescence minus one (FMO) controls for CD159a and CD4 to allow for accurate gating of GFP, PE, and PerCP/Cy5.5 fluorescence. Cells from the spleen and bronchial lymph nodes were mechanically isolated from freshly excised tissues using 18-gauge needles in cold RPMI and passed through a 100-μm cell strainer. Spleen cells were lysed using RBC lysis buffer. 10^6^ cells were resuspended in 100 μl of PBS 2% FBS solution for antibody staining using the same antibody cocktail as the PBMCs.

### Serum stability assay

3 mL blood was collected from the vein into a serum collection tube. The blood was placed at room temperature for 15 min. The coagulated blood was spun at 1000 x g for 15 min at 4°C. EVs were spiked into the serum at a dose similar to that achieved in the *in vivo* experiments, scaled down from an estimated 500 ml of macaque plasma to 50 μl serum. Thus, 4.7μL palmGRET EVs (dose 1:7.0E+06 EVs, dose 2:4.0E+07 EVs, dose: 3 2.0E+08 EVs, dose 4:9.0E+08 EVs) were spiked into 50 μl macaque serum (n= 2), and incubated for 0, 1,4 and 24 h at 37°C. EV stability was measured by nanoluciferase assay as described below.

### Nanoluciferase assays

Purified EV samples and SEC fractions were diluted 20-fold in PBS and were loaded into a white plastic 96 well plate, 50 μl per well, in duplicates. Biofluid samples, PBMC lysates and tissue homogenates, were loaded undiluted at 50 μl per well in duplicates. Nano-Glo substrate (furimazine, Promega, Madison, WI) was diluted 1:50 in assay buffer according to the manufacturer’s instructions. 50 μl diluted Nano-Glo reagent was added per well, and bioluminescence was measured immediately on a Fluoroskan Ascent plate reader (software v6.2) in bioluminescence mode, integration time 20 ms.

### IgG and cytokine measurements

Total IgG levels were measured in macaque plasma using a human IgG ELISA kit (Abcam, ab195215), which is cross-reactive with macaque IgG. Plasma samples, collected 24 hours after subsequent EV administrations, were diluted 70,000x in sample diluent, and IgG levels were measured following the manufacturer’s instructions. The plate was measured on a BioRad plate reader at 450 nm. Cytokine levels in macaque plasma were measured using the LEGENDplex NHP Inflammation Panel in filter plates (BioLegend #740332). Plasma samples were diluted 1:4 in assay buffer. Cytokine levels were measured following manufacturer’s instructions using a BD LSR Fortessa flow cytometer.

### Statistics and EV half-life calculation

Statistical differences were determined by one-way ANOVA with Tukey’s post-hoc test in GraphPad Prism 9.1, differences with p < 0.05 were considered to be statistically significant. EV half-life in biofluids was determined by linear regression of the nanoluciferase signal versus time on a log-lin chart. EV half-life was calculated from the slope of the regression line: t_1/2_ = log(2) / slope

## Availability of protocols

Procedural details have been submitted to the EV-TRACK knowledgebase and are available under EV-TRACK ID: EV210210^80^. High resolution flow cytometry experiments were performed following the MIFlowCyt-EV guidelines^40^.

## Author contributions

Drs. Tom Driedonks, Kelly Pate, and Kenneth Witwer conceived the research and designed experiments. Tom Driedonks, Linglei Jiang, Zheng Han, Guanshu Liu, Erin Shirk, Olesia Gololobova, Lyle Nyberg, Liliia Paniushkina, Marta Garcia-Contreras, Gabriela Lima and Barbara Smith performed experiments. Bess Carlson, Suzanne Queen, Zheng Han, Guanshu Liu, Kayla Schonvisky, Natalie Castell, Mitchell Stover, Selena Guerrero-Martin, Riley Richardson, Jessica Izzi, Eric Hutchinson and Kelly Pate performed animal experiments. Charles Lai designed and provided the palmGRET reporter construct. Tom Driedonks and Kenneth Witwer wrote the manuscript. All authors reviewed and revised the manuscript.

## Acknowledgements

The authors thank W. Zhang and T. Nilles for assistance with the imaging flow cytometry measurements.

---

**Supplemental Figure 1.**
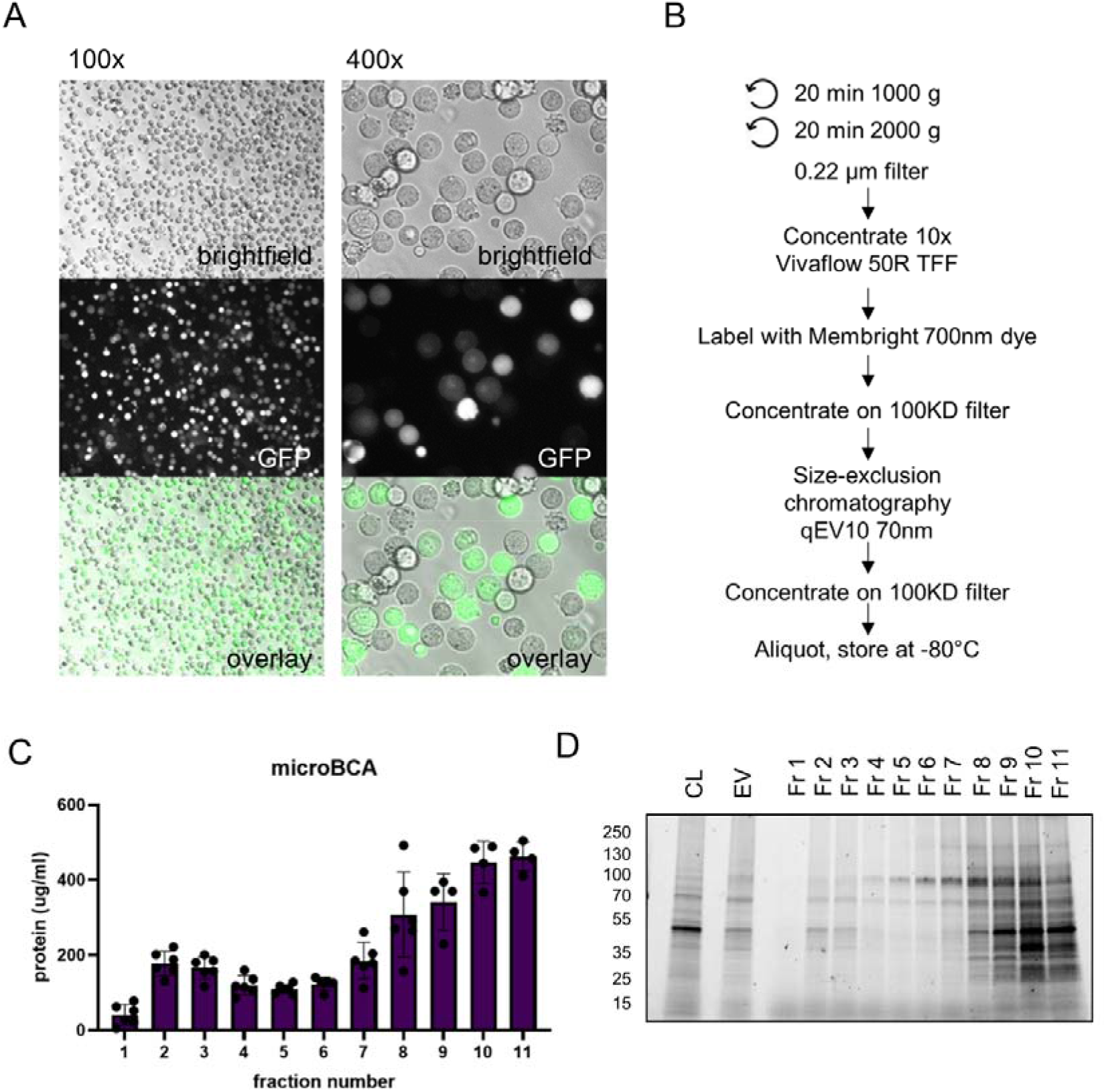
EV production and protein quantification. A) Microscopic images of Expi293F cells at harvest, three days after transfection. B) Schematic EV processing workflow. C) EVs produced by Expi293F cells were concentrated by TFF and ultrafiltration, followed by size-exclusion chromatography (SEC). Protein concentration on SEC fractions was determined by microBCA. Data from n=6 EV batches are shown. D) Stain-free imaging of SDS-PAGE on equal volumes of SEC fractions 1 – 11, pooled EVs (fr. 1-4) and cell lysate (CL). Image is representative of n=6 EV batches.

**Supplemental Table 1.** EV production and characterization parameters.

| EV batch nr | 1 | 2 | 3 | 4 | 5 | 6 |  | average | SD |
| --- | --- | --- | --- | --- | --- | --- | --- | --- | --- |
| cell dens harvest (c/ml) | 3.42E+06 | 4.50E+06 | 4.80E+06 | 3.00E+06 | 5.10E+06 | 5.70E+06 |  | 4.42E+06 | 1.03E+06 |
| % cell viability | 93.00% | 90% | 82% | 87% | 89% | 86% |  | 87.83% | 3.76% |
| EV protein conc (ug/ml) | 99 | 140.7 | 305.3 | 157.8 | 234 | 388.9 |  | 220.95 | 110.27 |
| EV Nluc in (RLU / 50 µl) | 1.44E+06 | 2.20E+07 | 6.27E+07 | 6.58E+07 | 5.68E+07 | 5.28E+07 |  | 4.4E+07 | 2.59E+07 |
| EV conc (p/ml) | 1.60E+11 | 4.40E+11 | 6.00E+11 | 4.50E+11 | 5.70E+11 | 7.60E+11 |  | 5.0E+11 | 2.02E+11 |
| particle/protein ratio | 1.62E+09 | 3.13E+09 | 1.97E+09 | 2.85E+09 | 2.44E+09 | 1.95E+09 |  | 2.3E+09 | 5.84E+08 |

**Supplemental Figure 2.**
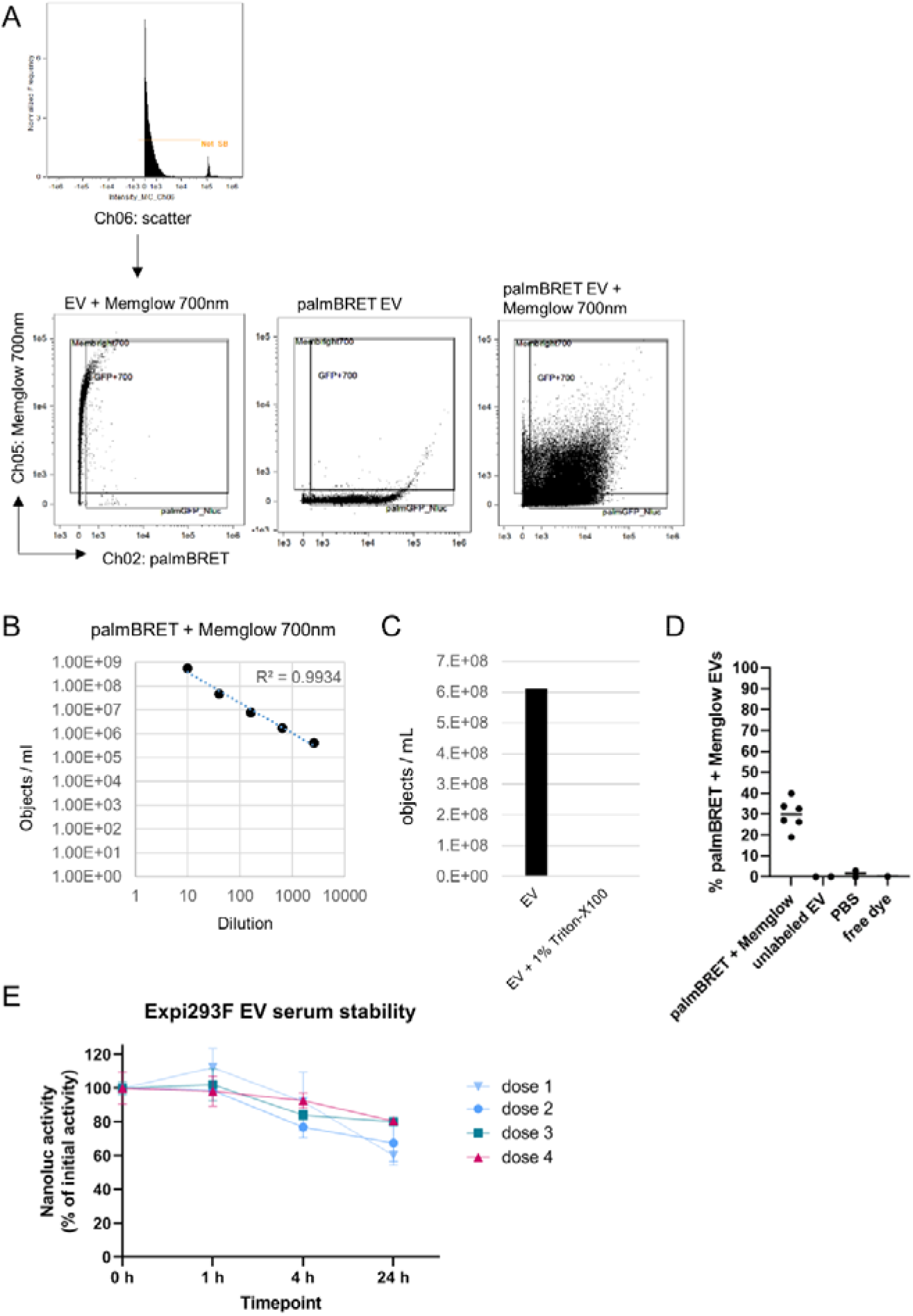
Characterization of palmGRET EVs by imaging flow cytometry. Different EV batches were analyzed by imaging flow cytometry. Single-stain controls, free dye control, and PBS were included for gating and determination of background signal. A) Gating strategy. Speed-beads were gated out, after which MemGlow and GFP gates were set based on Expi293 EV labelled only with MemGlow (left), and palmGRET-EV not labelled with MemGlow (middle). B) Dilution series of palmGRET-MemGlow EV shows the linearity of the measuring range. C) Addition of 1% Triton results in loss of palmGRET-MemGlow double positive objects. D) Percentage of palmGRET+MemGlow positive EVs in n=6 different batches, compared to unlabeled EV, PBS, and free dye (all measured twice). E) EV stability in macaque serum was measured by spiking macaque serum with EVs at doses equivalent to those achieved in the *in vivo* experiments. EV signal was measured by Nano-Glo assay after incubation for 1,4, or 24h at 37°C, expressed as % of the initial dose. N=2 measurements are shown.

**Supplemental Figure 3.**
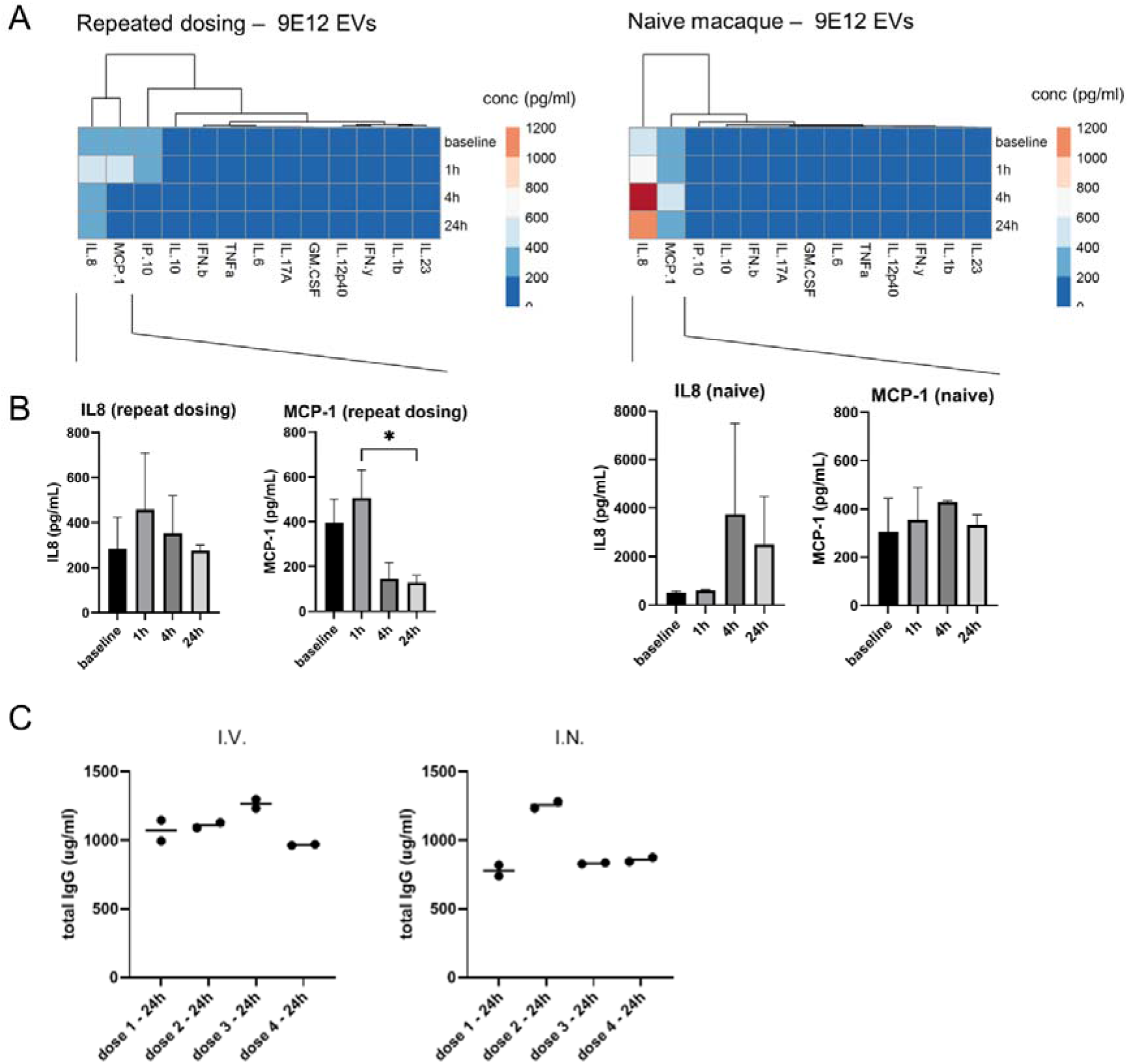
EV administration does not result in immune response. A) LegendPlex measurement of cytokine levels in macaque plasma collected before, and at 1, 4 and 24 h after IV administration of 9E12 EVs. Heatmaps are of cytokine levels in a subject previously dosed repeatedly with increasing doses (left) and in an EV-naïve subject (right) N=2 technical replicates. B) Bar graphs of the IL8 and MCP-1 levels from the measurements in A. * p < 0.05, determined by one-way ANOVA with Tukey’s post hoc test. C) Total IgG levels were determined by ELISA in macaque plasma collected 24h after each IV and IN dose. N=2 technical replicates.

**Supplemental Figure 4.**
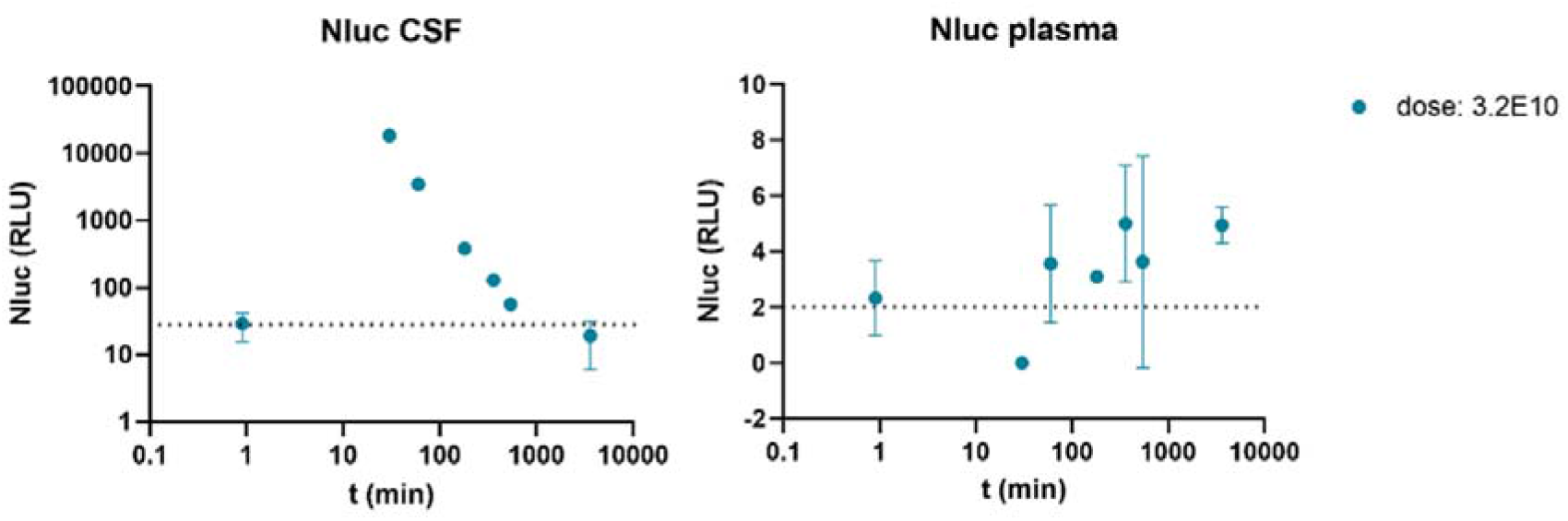
Intrathecal injection of EVs leads to rapid clearance from CSF. 3E10 palmGRET EVs were injected intrathecally into a macaque. CSF (left) and plasma (right) were sampled before and 0.5, 1, 3, 6, and 24 h after EV administration. EVs were detected in CSF and plasma by Nano-Glo assay. Error bars indicate assay variability.

**Supplemental Figure 5.**
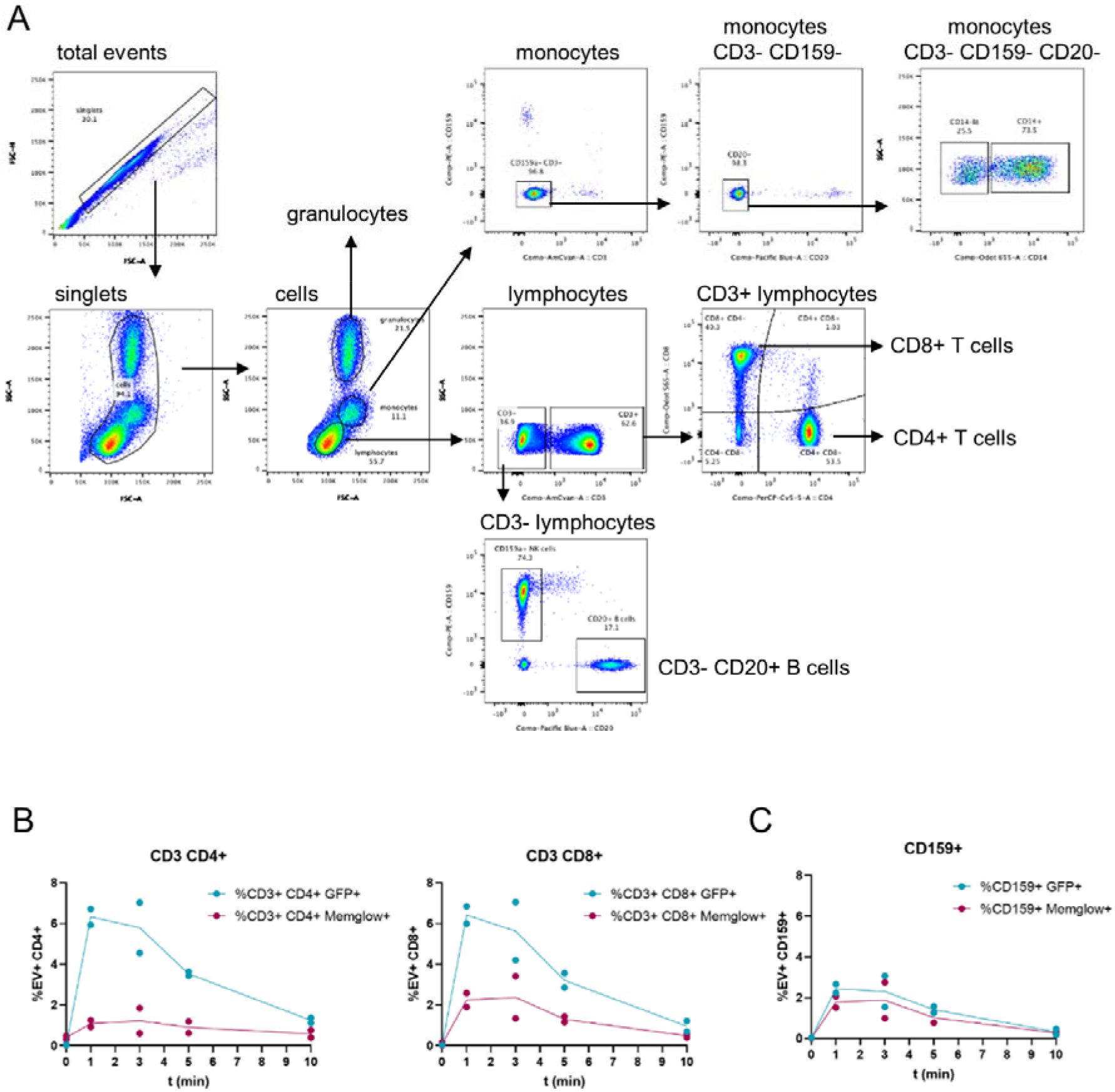
Flow cytometry of PBMCs in whole blood. A) Overall gating strategy. First, single cells were gated from total events. Monocyte, granulocyte and lymphocyte subsets were identified based on FSC / SSC profiles. Monocytes were identified by subsequently gating for CD3-CD159-cells, followed by gating for CD20-cells. Lymphocytes were divided into CD3+ cells, which contained both CD8+ and CD4+ T cells, and CD3-cells, including CD3-CD20+ B cells and CD3-CD159+ NK cells. B) Quantification of GFP+ and MemGlow+ CD4+ and CD8+ T cells, as a percentage of the total CD4+ and CD8+ T cells, respectively. Data from n=2 intravenous EV administrations are shown. C) Quantification of GFP+ and MemGlow+ positive CD3-CD159+ NK cells as a percentage of the total NK cells. Data from n=2 intravenous EV administrations are shown.

**Supplemental Figure 6.**
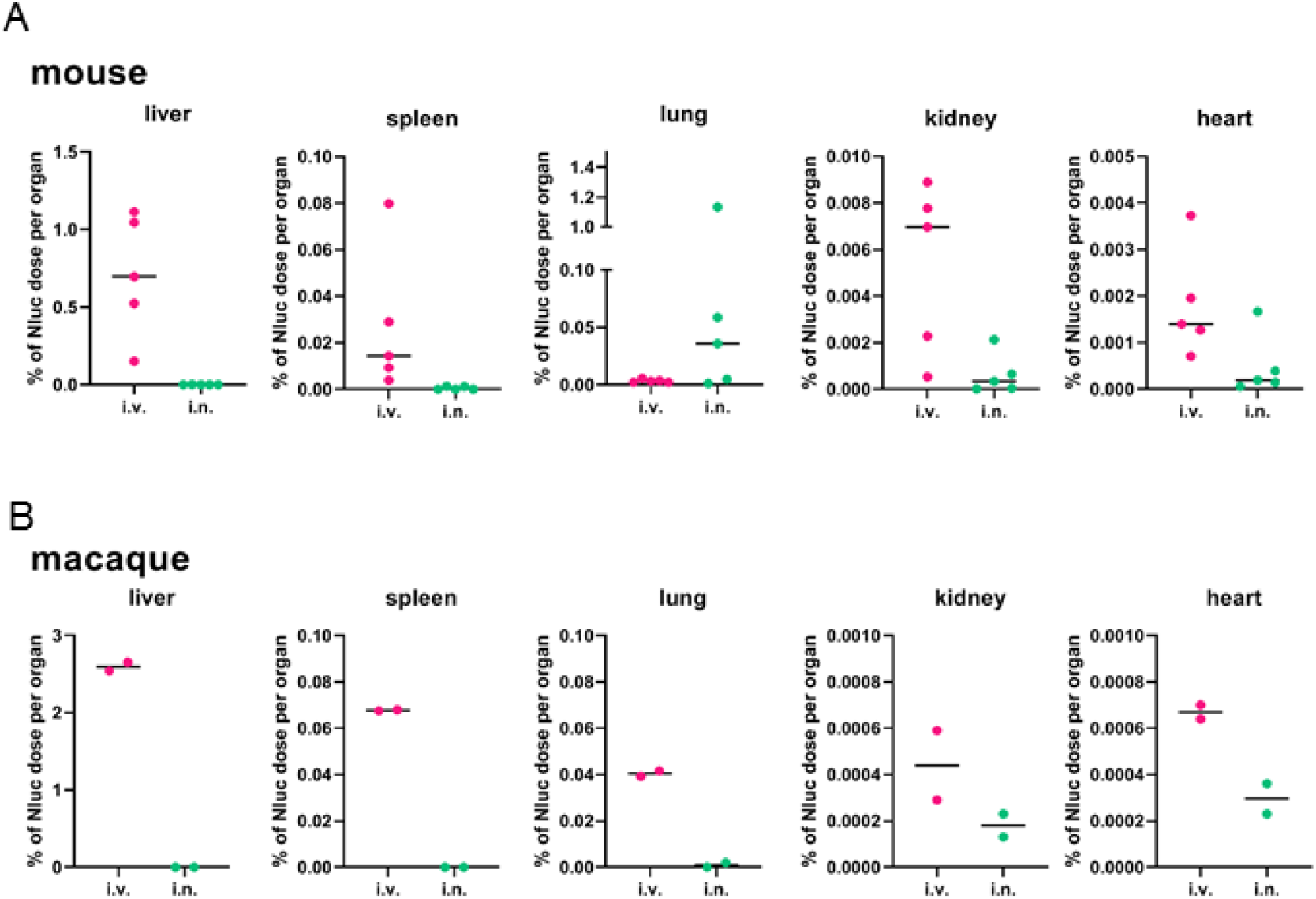
EV uptake by different organs in mice and macaques, as % of injected dose. A) EV uptake in mouse tissues was determined by Nano-Glo assay on tissue homogenates, as in Figure 4D. Nluc signal was expressed as % of the injected dose per whole organ. Data from n=5 animals are shown. B) EV uptake in macaque tissues was determined by Nano-Glo assay, as in Figure 5A. Nluc signal was expressed as % of the injected dose per whole organ. Data from 1 animal per group is shown, dots indicate replicate measurements of the same sample.

## Table of contents graphical abstract

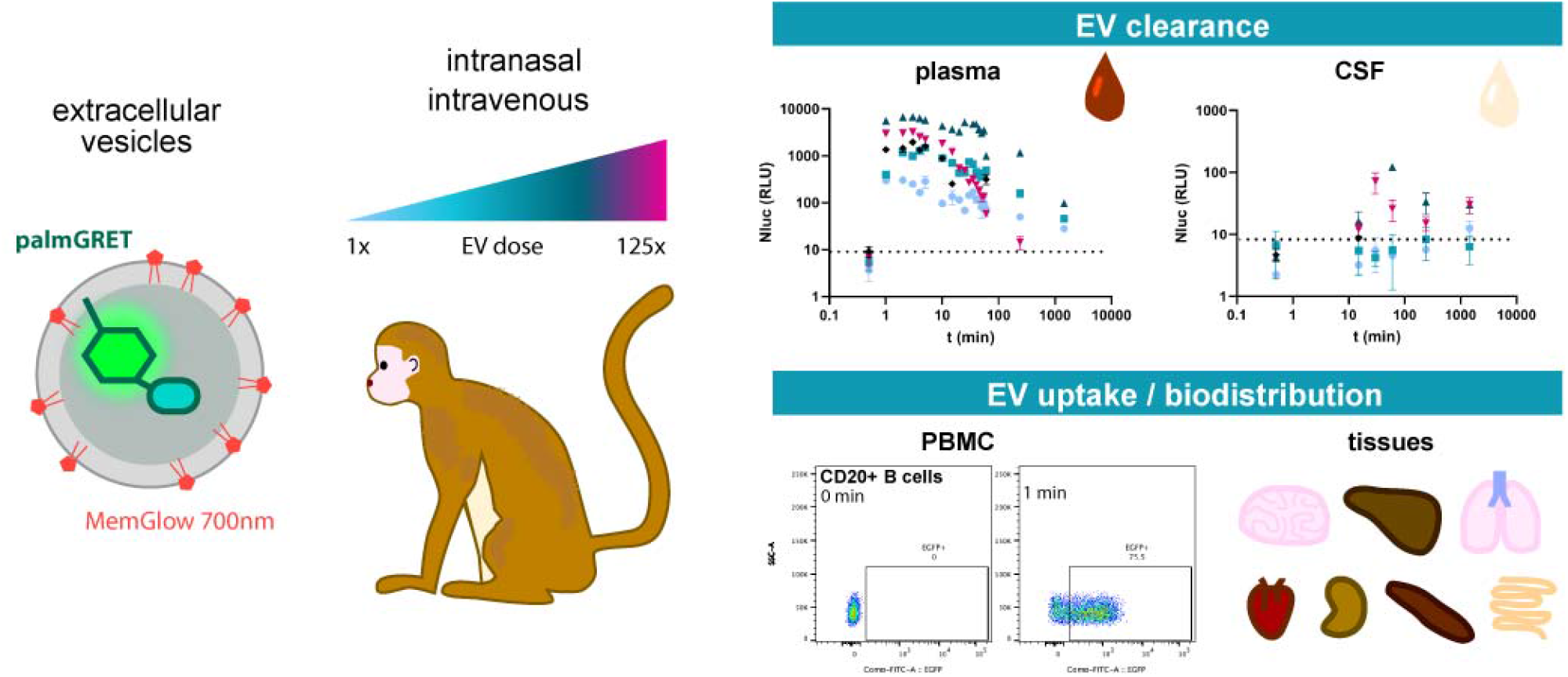

